# Specialized outputs and behavioral contributions of Purkinje cell subtypes

**DOI:** 10.64898/2026.05.20.726621

**Authors:** Shuting Wu, Chih-Chun Lin, Joon-Hyuk Lee, Kyoungdoo Hwang, Cameron A. Gayoso, Kanupriya Chawla, Naomi B. Johnston, Victor I. Ferrando, Wade G. Regehr

## Abstract

Purkinje cells (PCs), the sole outputs of the cerebellar cortex, transform granule cell firing patterns into appropriate outputs to drive learning and behavior. Two major PC classes (typically defined by *Aldoc* expression) can be further subdivided into nine molecularly distinct PC subtypes, suggesting that each subtype might be specialized for distinct categories of cerebellar processing. This has been difficult to test due to limited tools to target PC subtypes. We therefore developed intersectional tools to target PC subtypes, specifically Kcng4+ PCs (primarily Aldoc-) and Gpr176+ PCs (Aldoc1). We mapped their distributions within the cerebellar cortex and quantitatively characterized their outputs onto different types of cerebellar nuclei (CbN) neurons. Although projections by PC subtypes defined by Kcng4 and Gpr176 are regionally segregated within the CbN, each PC subtype synapses onto many types of CbN neurons, and individual CbN neurons often receive convergent inputs from multiple PC subtypes. We also selectively silenced PC subtype outputs and found that silencing Kcng4+ PC outputs impaired motor behaviors while sparing emotional and social behaviors, whereas silencing Gpr176+ PC outputs selectively increased exploratory behavior and reduced anxiety without affecting motor and social behaviors. These findings demonstrate that molecularly defined PC subtypes differentially regulate specific behaviors and establish a versatile framework for uncovering how cerebellar circuits are specialized to control diverse behaviors.

## Introduction

The cerebellum plays an essential role in a wide range of behaviors, including motor coordination, balance, posture, motor learning, cognitive function, language processing, reward processing, attention, affective behaviors, social behaviors, and emotional regulation ^1–8^. These diverse functions usually involve distinct regions of the cerebellar cortex ^9–13^, and there is growing recognition that these regions have specialized cell types and circuitry ^14,15^. Recent attention has focused on specializations of Purkinje cells (PCs)—the sole outputs of the cerebellar cortex ^15–17^. Mossy fiber inputs from diverse brain areas activate granule cells (GrCs) that then excite PCs, and learning is thought to rely to a large extent on GrC-PC synaptic plasticity ^14,18^. PCs project primarily to cerebellar nuclei (CbN), which in turn relay cerebellar outputs to the rest of the brain. Different types of excitatory neurons in the CbN project to distinct downstream regions that are implicated in different behaviors ^19–25^. These findings suggest that PC specializations have the potential to profoundly influence cerebellar processing in ways that are tailored to particular behaviors.

The observation that PCs are divided into two classes of cells based on differential expression of many different genes, including Aldoc (Aldolase C; also known as Zebrin II), established that PCs do not have uniform properties ^26–29^. Aldoc+ and Aldoc-PCs are present in alternating parasagittal stripes across the cerebellar cortex. Aldoc-PCs fire at higher frequencies ^30^ and are more susceptible to degeneration ^31^. Long-term plasticity at GrC-PC synapses also differs for Aldoc+ and Aldoc-PCs ^32–36^. Although Aldoc+ and Aldoc-PCs preferentially project to distinct regions of the CbN ^37–41^, the extent to which they selectively target different types of CbN neurons and have segregated pathways is unclear. Their different spatial distributions suggest distinct functional roles: Aldoc-PCs are preferentially located in regions associated with motor behaviors such as the anterior vermis, whereas Aldoc+ PCs are more prominent in regions implicated in vestibular, cognitive and affective functions including the posterior vermis, Crus I, flocculus and paraflocculus ^4,9,11,42^. However, these regions usually contain a mixture of Aldoc+ and Aldoc-PCs ^15,41^. Thus, despite well-established physiological and anatomical differences between Aldoc+ and Aldoc-PCs, it remains unclear whether they differentially regulate specific behaviors.

Single-nuclei RNA sequencing (snRNA-seq) studies of the cerebellar cortex revealed seven subtypes of Aldoc+ and two subtypes of Aldoc-PCs with distinct molecular profiles ^15,17,43^. This additional diversity suggests that PCs can influence cerebellar processing with greater flexibility than had been appreciated. Each subtype is preferentially enriched in specific lobules, and most lobules contain multiple PC subtypes ^15^. The identification of subtype-specific markers along with *in situ* hybridization (ISH) studies allow more precise mapping of the distribution of PC subtypes; for example, *Gpr176* can be used to label Aldoc1 PCs ^15,44^. However, investigating the specializations of these molecularly defined PC subtypes has been hindered by the lack of tools to selectively target PC subtypes.

Here, we developed the tools required to implement an intersectional strategy to selectively target PC subtypes and determine their circuit properties and behavioral roles. We used this approach to label specific PC subtypes, map their distributions within the cerebellar cortex, and quantitatively characterize their outputs onto different types of CbN neurons. We also selectively silenced the outputs of PC subtypes to assess their differential contributions to behaviors. We found that silencing Kcng4+ PCs (that consist primarily of Aldoc-PCs) disrupted many motor behaviors while sparing emotional and social behaviors. In contrast, silencing Gpr176+ PCs (Aldoc1 PCs) selectively increased exploratory behavior and reduced anxiety while sparing motor and social behaviors. Together, these findings demonstrate that distinct PC subtypes form specialized cerebellar pathways to differentially regulate specific behaviors, shifting our view of cerebellar function from a solely regional organization to a framework that integrates regional organization with specialized circuits and opening the door to a broader understanding of how cerebellar circuits shape diverse behaviors.

## Results

### An intersectional approach to target PC subtypes

To study the circuit and function specializations of PC subtypes, it is vital to selectively target and manipulate these subtypes. Several transgenic lines, including Aldoc-Venus, EAAT4-GFP, and Kcng4-Cre, have been used to label different PC subtypes ^41,45,46^. However, these transgenic lines lack PC-specificity and often label other types of neurons in the cerebellum and numerous other brain regions. An intersectional strategy based on Cre and Flp lines is widely used to achieve cell type specificity ^47^. Although PCs can be selectively targeted with a Pcp2^Cre^ line ^48^, the limited number of suitable Flp lines limits the use of Pcp2^Cre^ in an intersectional approach. Therefore, we generated a Pcp2^Flp^ line to selectively target PCs (**Fig. S1A**), which was chosen based on the PC selectivity of the Pcp2^Cre^ line ^48^. When we crossed it with a Flp reporter line, RC::FLTG ^49^, the restricted expression of tdTomato (tdT) in PCs established the selectivity and suitability of the Pcp2^Flp^ line (**Fig. 1A**).

**Figure 1.**
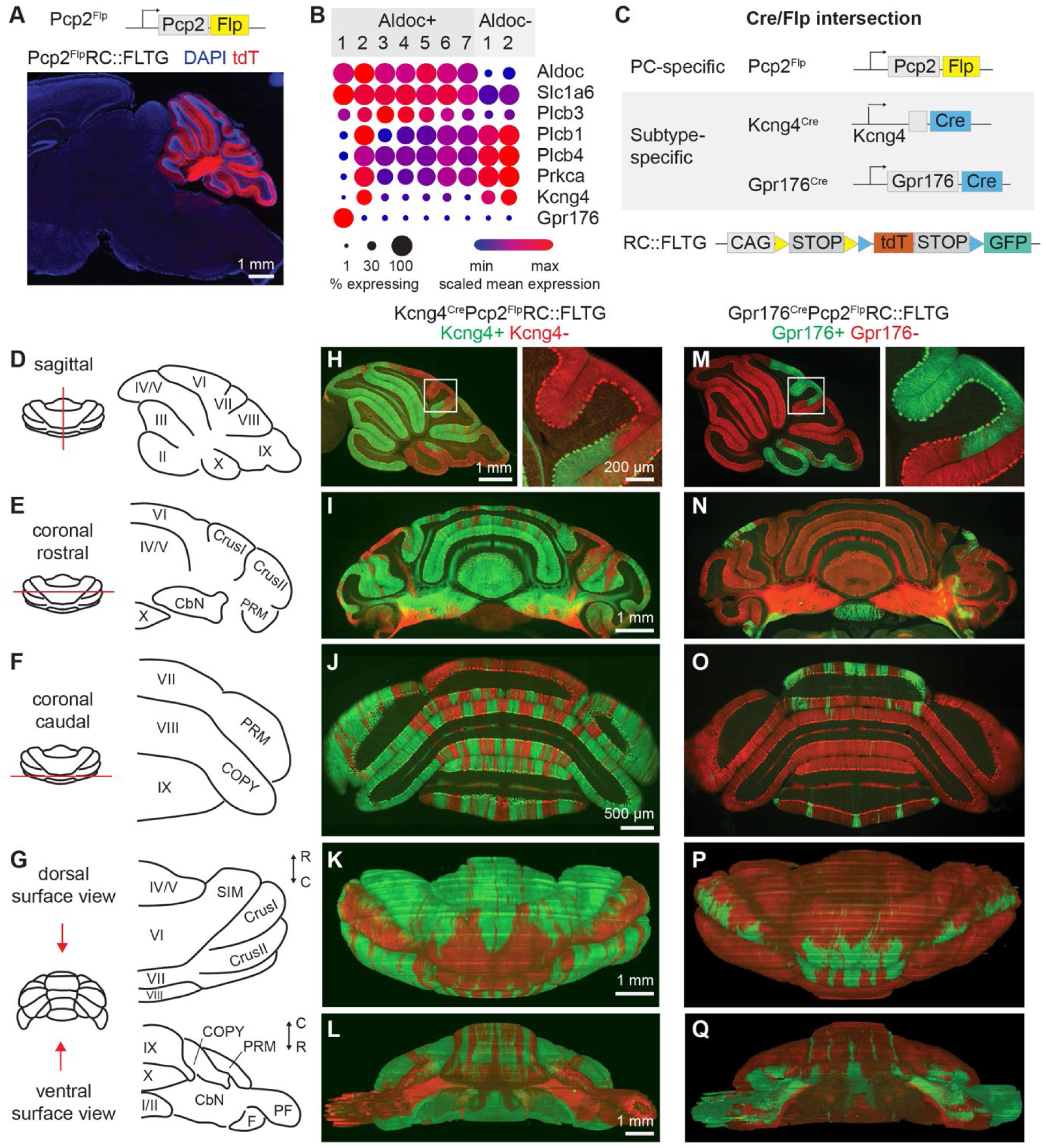
An intersectional approach to target PC subtypes. **A.** A sagittal brain section in a Pcp2^Flp^RC::FLTG mouse shows tdT labeling is restricted to PCs. **B.** Dot plots show expression profiles for the indicated genes in 9 PC subtypes ^15^. **C.** Diagram of Cre/Flp intersection strategy using a PC-specific Flp driver line, subtype-specific Cre driver lines, and a Cre/Flp-dependent reporter line. **D.** A sagittal section (red line) across the midline of cerebellum which is shown on the right. **E.** A coronal section across the rostral cerebellum which is shown on the right. **F.** Same as **E** but for caudal cerebellum. **G.** A 3D reconstruction of the cerebellum from serial coronal sections, with dorsal and ventral surface view shown on the right. **H-L**. Fluorescence images of the Kcng4^Cre^Pcp2^Flp^RC::FLTG mice in which Kcng4+ PCs are labeled with GFP (*green*) and Kcng4-PCs are labeled with tdT (*red*). **H.** A sagittal section in a Kcng4^Cre^Pcp2^Flp^RC::FLTG mouse corresponding to **D**. *Inset*, magnified image of the outlined area. **I.** A coronal section in the rostral cerebellum of a Kcng4^Cre^Pcp2^Flp^RC::FLTG mouse corresponding to **E**. **J.** Same as **I** but for caudal cerebellum corresponding to **F**. **K.** A dorsal surface view of the cerebellum of a Kcng4^Cre^Pcp2^Flp^RC::FLTG mouse corresponding to **G** (*top*). **L.** Same as **K** but for a ventral surface view corresponding to **G** (*bottom*). **M-Q**. same as **H-L**, but for the Gpr176^Cre^Pcp2^Flp^RC::FLTG mice in which Gpr176+ PCs are labeled with GFP (*green*) and Gpr176-PCs are labeled with tdT (*red*). Abbreviations: SIM, Simple lobule; PRM, Paramedian lobule; COPY, Copula pyramidis; PF, Paraflocculus; F, Flocculus.

We began our search for appropriate genes and Cre lines for PC subtypes by focusing on the widely studied Aldoc+ and Aldoc-PCs. Numerous markers have been identified to be differentially expressed in these two populations, including *Aldoc, Plcb3, Slc1a6, Plcb1, Plcb4, Prkca,* and *Kcng4* ^26,28,46,50–53^. Differential expression of all these genes was evident in previous snRNA-seq studies ^15^, but in most cases the expression is not perfectly restricted to a single PC subtype (**Fig. 1B**). Among Aldoc+ population markers, both *Aldoc* and *Slc1a6* are also expressed at low levels in Aldoc-PCs, whereas *Plcb3* is expressed at various levels in different Aldoc+ subtypes. Among Aldoc-markers, *Kcng4* (which encodes the potassium voltage-gated channel Kv6.4) is the most promising candidate because it is expressed primarily in Aldoc-PCs and Aldoc2 PCs, whereas *Plcb1*, *Plcb4*, and *Prkca* are also present at reasonably high levels in most Aldoc+ subtypes. Aldoc2 PCs are a small population of PCs (less than 0.5% of all PCs) that are specifically found in the flocculus (F) and paraflocculus (PF) ^15^, so *Kcng4* preferentially labels Aldoc-PCs in the vermis and hemisphere of the cerebellum. A Kcng4^Cre^ line has recently been used to preferentially target Aldoc-PCs ^46,54^. We therefore acquired this line and confirmed that in Kcng4^Cre^Ai14 mice, stripes of PCs were labeled in an Aldoc-pattern throughout the cerebellar cortex (**Fig. S1C**). Extensive labeling was also seen in a subset of molecular layer interneurons (MLIs) in the cerebellum and numerous other brain regions (**Fig. S1C**), indicating that an intersectional strategy is needed to selectively target Kcng4+ PCs.

We also wanted to target a specific PC subtype within the 9 subtypes identified in our previous snRNA-seq study (7 Aldoc+ subtypes and 2 Aldoc-subtypes) ^15^. We are particularly interested in the Aldoc1 subtype, because it has the most distinct molecular profile with many candidate marker genes and is found in regions that are implicated in nonmotor behaviors ^55,56^. One of the candidate genes is *Gpr176*, which encodes an orphan G-protein-coupled receptor (**Fig. 1B**). ISH shows that *Gpr176* is expressed in distinct groups of PCs in lobule VI/VII, IX, X, F and PF ^15,44^. A suitable Cre line was not available for Aldoc1 PCs, so we made a Gpr176^Cre^ line (**Fig. S1B**). In Gpr176^Cre^Ai14 mice, the tdT labeling pattern is observed in a subset of PCs in regions consistent with ISH studies, as well as a subset of MLIs and numerous other brain regions (**Fig. S1D**).

We then crossed the respective Cre lines with Pcp2^Flp^ and RC::FLTG, in which Flp+Cre+ cells express GFP and Flp+Cre-cells express tdT (**Fig. 1C**). This will specifically label PCs either with GFP or tdT while sparing all other neurons. We characterized the cell labeling patterns in these triple transgenic mice using sagittal sections (**Fig. 1D**), coronal sections (**Fig. 1E and 1F**), and 3D reconstructions based on 50 μm-thick serial coronal sections (**Fig. 1G**). In sagittal sections of Kcng4^Cre^Pcp2^Flp^RC::FLTG mice, PC somata, dendrites, and axons were labeled with either GFP or tdT, while all other neurons were unlabeled (**Fig. 1H**). Coronal sections revealed parasagittal stripes of GFP+ and tdT+ PCs throughout the cerebellum (**Fig. 1I and 1J**; for F and PF, see **Fig. S2A**) and their axon projections in the CbN (**Fig. 1I**). In the 3D cerebellar reconstructions, Kcng4^Cre^Pcp2^Flp^RC::FLTG mice label stripes across multiple lobules of the cerebellum, with Kcng4+ PCs mainly found in anterior vermis and hemisphere, and Kcng4-PCs found in central and posterior vermis, Crus I, and P/PF (**Fig. 1K and 1L, Supplementary video 1**), which is reminiscent of the Aldoc pattern described previously ^29,37,41^.

The labeling pattern of Gpr176^Cre^Pcp2^Flp^RC::FLTG mice was very different. GFP labeling was restricted to small groups of PCs in lobule VI, VII, IX and X in the vermis, Crus I in the hemisphere, and F/PF (**Fig. 1M-Q; Fig. S2B, Supplementary video 2**), which appears to be a subset of the Kcng4-PCs when compared to the Kcng4^Cre^Pcp2^Flp^RC::FLTG mice (**Fig. 1H-L**). This pattern is consistent with the *Gpr176* ISH and the lobule enrichment pattern from previous RNA-seq studies ^15,44^. GFP+ PCs are either in the form of stripes (lobule VI, VII, IX, X, **Fig. 1O-Q**), or patches on the surface of the cerebellar cortex (Crus I, **Fig. 1N and 1P**). There is almost no GFP labeling in the anterior vermis (**Fig. 1N and 1P**) and extensive labeling in the PF (**Fig. 1Q; Fig. S2B**).

The 3D reconstruction of the cerebellum only reveals the pattern of PCs on the surface but not the subsurface PCs (**Fig. 2A**). To better visualize the distribution of PC subtypes in the entire cerebellar cortex, we applied an unfolding algorithm to generate a cerebellar 2D map (**Fig. 2B; Fig. S3; Methods**). We developed an unfolding algorithm in Python with an easy-to-use interface that is readily adapted for other kinds of images (**Fig. S3; Methods**). A schematic of the 2D map is shown in **Fig. 2C**, with individual lobules segmented, and the surface area highlighted in grey (39% of the total Purkinje cell layer). This 2D map was generated from sagittal sections and provides a more realistic aspect ratio of cerebellar cortex than the commonly used 2D maps based on coronal or horizontal sections ^37,41^. We then generated 2D maps with the PC labeling pattern in the triple transgenic mice. In the Kcng4^Cre^Pcp2^Flp^RC::FLTG mice, Kcng4+ PCs are mainly found in the anterior vermis and hemisphere, while Kcng4-PCs are present in stripes in central and posterior vermis, Crus I, F and PF (**Fig. 2D**). Kcng4 labels 69% of all PCs, 67% in the vermis, 82% in the hemispheres, and 54% in the F and PF. These patterns are largely reminiscent of the Aldoc pattern described previously, although there are minor differences, especially in the posterior vermis and hemisphere, where there are more Kcng4+ PCs than Aldoc-PCs ^29,37,41^. For Gpr176^Cre^Pcp2^Flp^RC::FLTG mice, Gpr176+ PCs are mainly found in lobule VI, VII, X, and PF, and to a lesser degree in lobule IX, Crus I, and F (**Fig. 2E**). Gpr176+ PCs account for 9% of all PCs, 6.8% in the vermis, 2.5% in the hemispheres, and 44% in the flocculus and paraflocculus. Many Gpr176+ PCs are present on the surface; therefore, the 2D map gives a more accurate representation of the percentage than the 3D reconstructions (**Fig. 1P and 1Q**). Even though Gpr176+ PCs (Aldoc1 PCs) comprise a low percentage of PCs, they are found in cerebellar regions that are implicated in diverse motor and non-motor behaviors.

**Figure 2.**
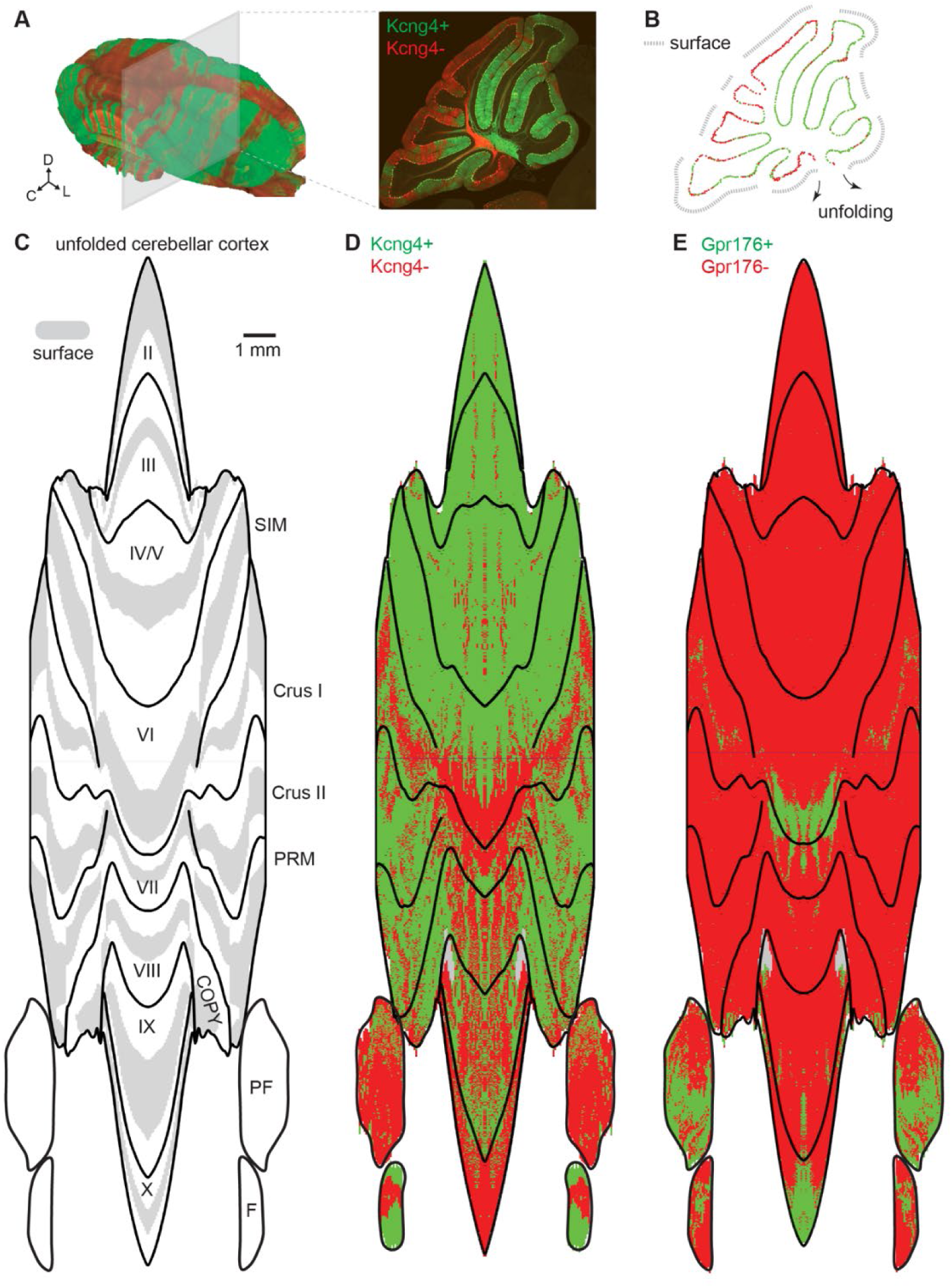
Unfolded maps of the cerebellar cortex. **A.** *Left*, a reconstruction of the cerebellum in the Kcng4^Cre^Pcp2^Flp^RC::FLTG mouse from serial coronal sections; *right*, a sagittal section showing details in the folded lobules. **B.** Cell segmentation of all the GFP+ and tdT+ PCs in the example sagittal section in **A**, with the grey dash line mark the surface area. To reveal the pattern of folded areas that are not visible in the reconstructed 3D cerebellum, we run a pipeline to unfold the continuous PCL in the whole cerebellar cortex (see **Fig. S3** and **Methods**). **C.** A 2D map showing an unfolded PCL of the whole cerebellar cortex, with surface area labeled in grey. **D.** A 2D map showing the distribution of Kcng4+ (green) and Kcng4-(red) PCs in the unfolded cerebellar cortex in a Kcng4^Cre^Pcp2^Flp^RC::FLTG mouse. The Image was mirrored from half of the cerebellum. **E.** Same as **D** but for Gpr176^Cre^Pcp2^Flp^RC::FLTG mouse. The Image was mirrored from half of the cerebellum.

### Differential firing rates of Aldoc1 and other Aldoc+ PCs

Previous studies have shown that Aldoc+ and Aldoc-PCs differ in many ways^32,57,58^, but it has not been possible to determine the properties of specific Aldoc+ subtypes. The firing rates of Aldoc-PCs are 1.5 to 2-fold higher than those of Aldoc+ PCs^30,59–61^. We examined whether the firing properties of Aldoc1 PCs differ from other Aldoc+ PC subtypes in lobule X of the cerebellar vermis, which contains only Aldoc+ PCs ^37,41^. Serial sections through vermal lobule X in Gpr176^Cre^Pcp2^Flp^RC::FLTG mice showed that the regional distributions of Gpr176+ (Aldoc1) PCs and Gpr176-PCs vary markedly with distance from the midline (**Fig. 3A**): in the most medial sections, almost all PCs are Gpr176+, whereas in more lateral sections Gpr176-PCs are restricted to the dorsal folium of lobule X and Gpr176+ PCs are found in the ventral folium. *Aldoc* ISH of comparable sections confirmed high *Aldoc* expression in all lobule X PCs (**Fig. 3B**). We used extracellular recordings in brain slices to assess the firing properties of lobule X PCs in the presence of ionotropic glutamate and GABA receptor antagonists. Gpr176+ (green) and Gpr176-(red) PCs were identified by fluorescence labelling in Gpr176^Cre^Pcp2^Flp^RC::FLTG mice. As described previously, many PCs fired regularly under these conditions (**Fig. 3C**), whereas other PCs displayed a triphasic firing pattern consisting of epochs of regular firing, bursts, and silent periods ^62^. Gpr176+ PCs generally fired at a lower rate than Gpr176-PCs (**Fig. 3C**), with a characteristic action potential waveform with smaller positive peak and slower repolarization (**Fig. 3D**). For regularly firing PCs, the average firing rate of Gpr176+ PCs was lower than Gpr176-PCs [33.7 ± 2.7 spikes/s (n=22, N=3, ± SEM) vs. 48.9 ± 4.2 spikes/s (n=18, N=3)], with a higher CV2 [0.049 ± 0.004 (n=22, N=3) vs. 0.032 ± 0.004 (n=18, N=3)] (**Fig. 3E**). For irregularly firing PCs, the average firing rate of Gpr176+ PCs was also lower than Gpr176-PCs [44.5 ± 1.2 spikes/s (n=14, N=3) vs. 94.0 ± 2.2 spikes/s (n=19, N=3)] but there was no significant difference in CV2 [0.24 ± 0.04 (n=14, N=3) vs. 0.192 ± 0.014 (n=19, N=3)] (**Fig. 3F**). These findings indicate specific Aldoc+ PC subtypes have specialized firing properties, with Aldoc1 PCs firing 31% slower for regular spiking cells and 53% slower for irregular firing cells than other Aldoc+ PCs in lobule X.

**Figure 3.**
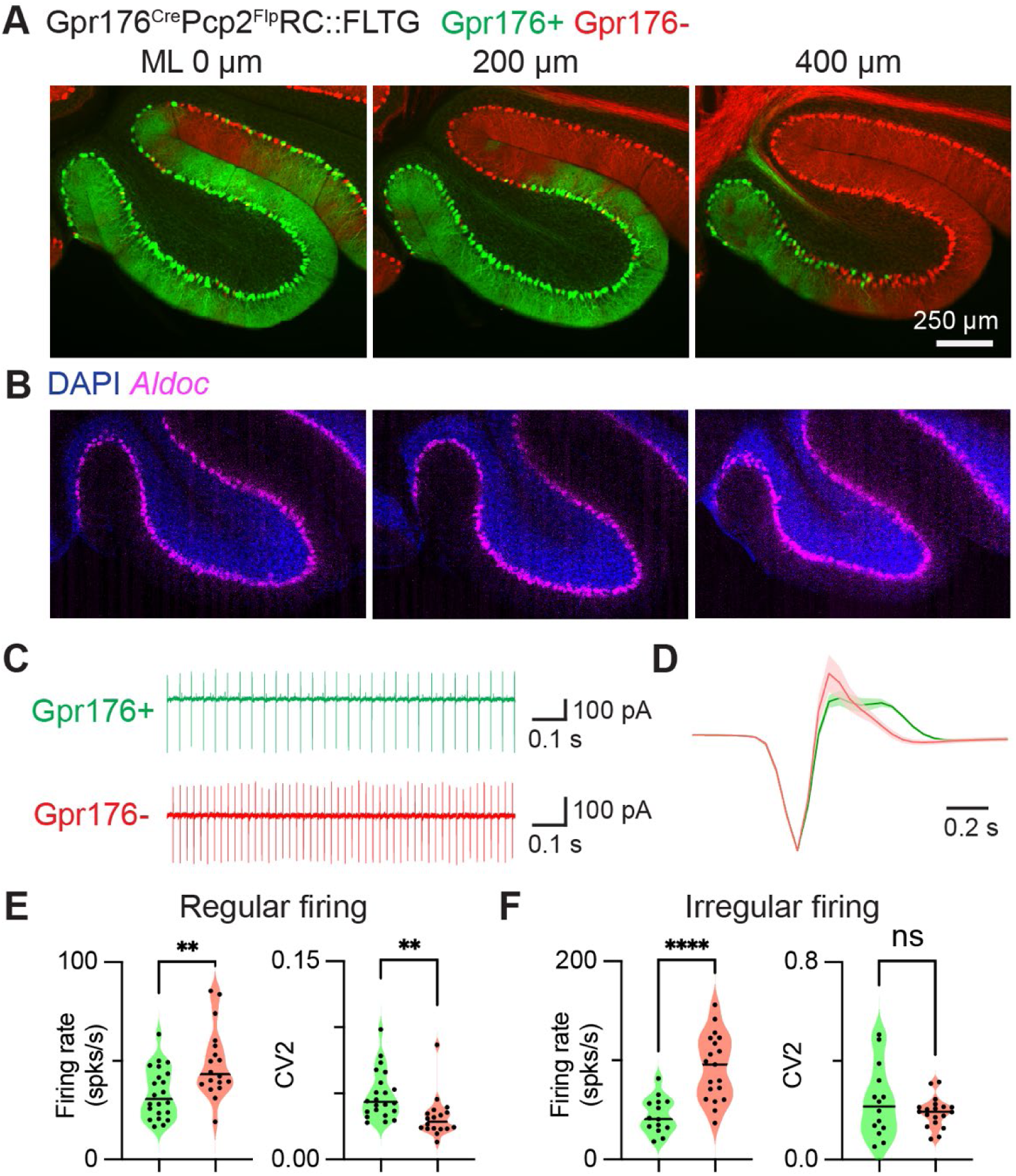
Firing properties of PC subtypes. **A.** Serial sections at different medial lateral (ML) locations across lobule X in a Gpr176^Cre^Pcp2^Flp^RC::FLTG mouse showing the varying distribution patterns of Gpr176+ (green) and Gpr176-PCs (red). **B.** *Aldoc* ISH in similar sections showing mot PCs in lobule X are Aldoc+. **C.** Representative traces of *in vitro* on-cell recordings for a Gpr176+ and a Gpr176-PC in lobule X guided by the fluorescent labeling in Gpr176^Cre^Pcp2^Flp^RC::FLTG mice. **D.** Representative action potential waveforms averaged from 6 Gpr176+ and 6 Gpr176-PCs with the biggest amplitudes in lobule X. **E.** Average firing rates and CV2 (local coefficient of variation) over a 3-mins period for the regularly firing Gpr176+ (n=22) and Gpr176-PCs (n=18). **F.** As in **E** but for irregularly firing Gpr176+ (n=14) and Gpr176-PCs (n=19).

### PC subtypes target distinct regions of the cerebellar nuclei and brainstem

Fluorescence labeling of PCs and their axons using this intersectional approach also allowed us to trace the outputs of PC subtypes to downstream regions. We primarily focused on the CbN, which are the main target of PCs and the main outputs of cerebellum. The CbN are comprised of three types of subnuclei bilaterally, the fastigial nucleus (FN), the interposed nucleus (IP), and the dentate nucleus (DN) (**Fig. 4A**). Previous studies characterized PC projections using Aldoc immunohistochemistry, fluorescence labeling in Aldoc-Venus mice, localized injections, and single PC reconstructions showed that Aldoc+ and Aldoc-PCs project to distinct regions of the CbN ^38,41,63^. We extended these studies by using an intersectional approach that labeled complementary groups of PCs with either GFP or tdT (**Fig. 1**).

**Figure 4.**
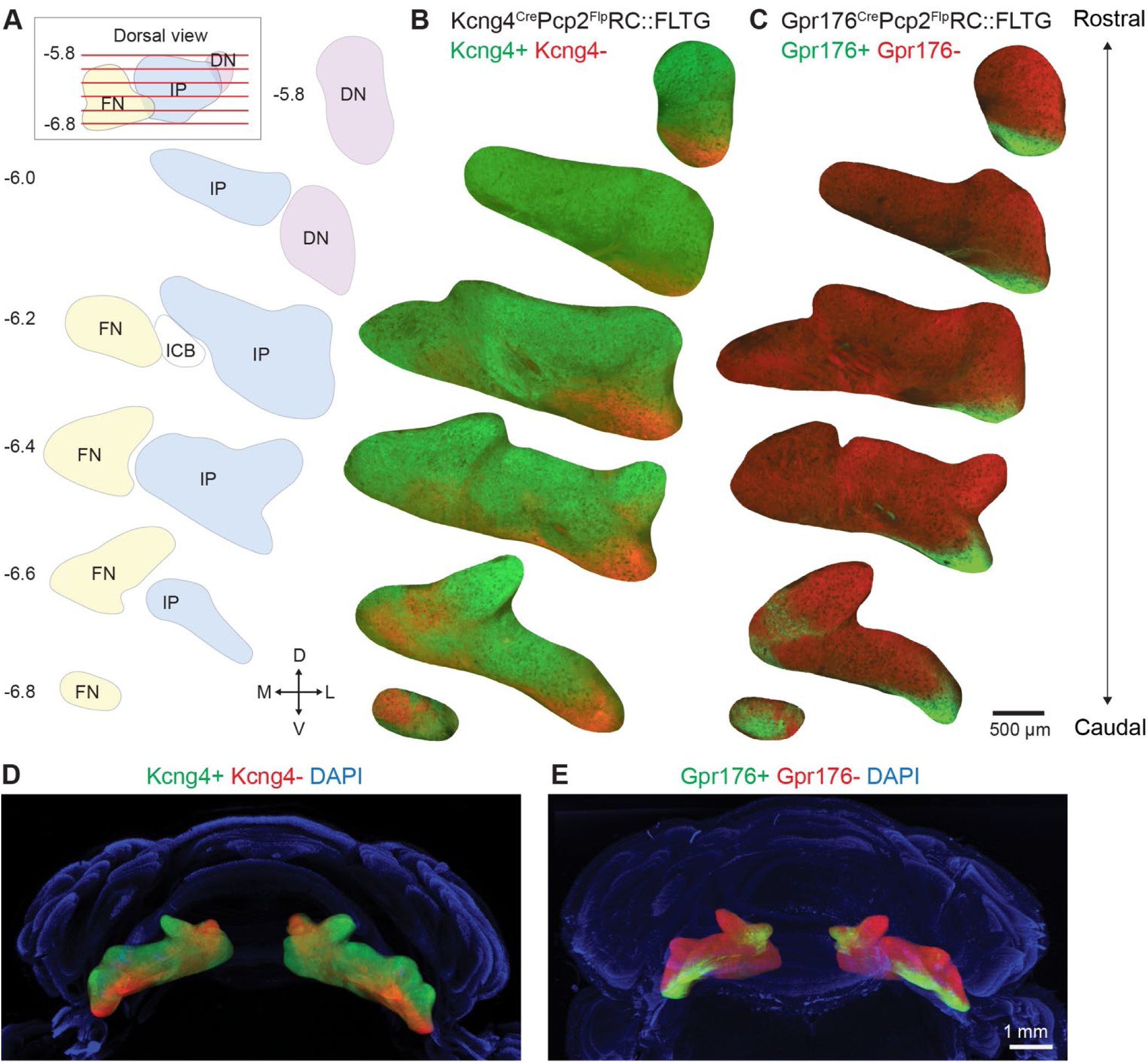
CbN projections by PC subtypes. **A.** Serial coronal sections of the right CbN with bregma positions indicated in the inset. **B.** Fluorescence images corresponding to the sections in **A** show the PC projections to CbN by Kcng4+ (green) and Kcng4-(red) PCs in a Kcng4^Cre^Pcp2^Flp^RC::FLTG mouse. Signal outside of the CbN were masked out. **C.** Same as **B** but for a Gpr176^Cre^Pcp2^Flp^RC::FLTG mouse, in which Gpr176+ PCs are labeled with GFP (green) and Gpr176-PCs are labeled with tdT (red). **D.** A caudal view of the 3D reconstruction of the cerebellum shows projections to CbN made by Kcng4+ (green) and Kcng4-(red) PCs in a Kcng4^Cre^Pcp2^Flp^RC::FLTG mouse. **E.** Same as **D** but for a Gpr176^Cre^Pcp2^Flp^RC::FLTG mouse.

This approach provides greater flexibility to target different PC subtypes and allows us to directly examine the degree of segregation and overlap of axonal signals from different PC subtypes. We characterized the PC subtype outputs using Kcng4^Cre^Pcp2^Flp^RC::FLTG and Gpr176^Cre^Pcp2^Flp^RC::FLTG mice and a series of coronal sections that span the whole CbN to visualize the projection patterns by PC subtypes (**Fig. 4B and 3C**). In Kcng4^Cre^Pcp2^Flp^RC::FLTG mice, Kcng4+ PCs (GFP) provide the majority of inputs to the CbN, whereas Kcng4-PCs (tdT) preferentially target small regions in the ventral DN, the ventral-lateral IP, and caudal-medial FN (**Fig. 4B**). Overall, Kcng4+ and Kcng4-PC projections are regionally segregated, with substantial overlap in transitional zones. In Gpr176^Cre^Pcp2^Flp^RC::FLTG mice, Gpr176+ PCs (GFP) label a subset of the regions innervated by Kcng4-PCs (**Fig. 4C**). Gpr176+ and Gpr176-PC axons show extensive overlap in caudal-medial FN. Together, these results indicate that PC subtypes defined by Kcng4^Cre^ and Gpr176^Cre^ provide largely segregated outputs to the CbN, with considerable overlap in transitional areas. In addition, we used 3D reconstructions of the cerebellum and masked out all regions except the CbN to better visualize PC projection patterns across the entire CbN for a Kcng4^Cre^Pcp2^Flp^RC::FLTG mouse and a Gpr176^Cre^Pcp2^Flp^RC::FLTG mouse (**Fig. 4D and 3E; Supplementary video 3** and **Supplementary video 4**). The labeling pattern in the CbN of the Kcng4^Cre^Pcp2^Flp^RC::FLTG mouse showed a dorsolateral and ventrocaudal separation, which is overall consistent with previous characterizations of PC projections using Aldoc immunohistochemistry and Aldoc-Venus mice with minor differences observed in the DN, consistent with the differences of PC labeling in the hemisphere ^37,41^.

PCs also provide direct outputs to numerous brainstem regions, including vestibular nuclei (VN) and the parabrachial nucleus (PBN) ^38,64–70^. Serial coronal sections across different brainstem regions in a Kcng4^Cre^Pcp2^Flp^RC::FLTG mouse showed that Kcng4+ PCs project to the superior vestibular nucleus (SUV), lateral vestibular nucleus (LAV), spinal vestibular nucleus (SPIV), and the ventral-lateral region of medial vestibular nucleus (MV), while Kcng4-PC axons project to the PBN, dorsal-medial MV, and nucleus prepositus (PRP) (**Fig. S4A-D**). In Gpr176^Cre^Pcp2^Flp^RC::FLTG mice, Gpr176+ PCs primarily project to dorsal-medial MV and PRP (**Fig. S4E-H**). Projection patterns are readily visualized in 3D reconstructions of Kcng4^Cre^Pcp2^Flp^RC::FLTG and Gpr176^Cre^Pcp2^Flp^RC::FLTG mice with fluorescence outside the brainstem masked out (**Fig. S4I and S4J; Supplementary video 5** and **Supplementary video 6**). In Kcng4^Cre^Pcp2^Flp^RC::FLTG mice, many Kcng4+ and Kcng4-PC axons exit the cerebellum via the inferior cerebellar peduncle (**Supplementary video 5**). Kcng4+ PC axons primarily project to nearby vestibular regions (**Fig. S4I; Supplementary video 5**), whereas Kcng4-PC axons initially extend caudally, then turn medially and rostrally before ultimately targeting medial brainstem regions, including the PRP and medial MV (**Fig. S4I; Supplementary video 5**). Gpr176+ PCs follow a trajectory similar to that of Kcng4-PC axons, projecting predominantly to medial brainstem regions (**Fig. S4J; Supplementary video 6**). Thus, different PC subtypes target distinct regions in the CbN and brainstem.

### PC subtypes target distinct neuron subtypes in the fastigial nucleus

The CbN contain multiple types of neurons that project to distinct downstream targets and regulate diverse behaviors ^20,21^. To gain insight into potential behavioral roles of different PC subtypes, we refined our characterization of PC output pathways by quantifying PC subtype inputs to individual CbN neurons, as shown for a Kcng4^Cre^Pcp2^Flp^RC::FLTG mouse and a caudal section of the FN (**Fig. 5**). Confocal grey-scale fluorescence images revealed dense axonal innervation from GFP+ PCs (Kcng4+, **Fig. 5A**) and tdT+ PCs (Kcng4-, **Fig. 5B**) within the FN, with pronounced regional specificity. tdT was predominantly localized to the caudal-medial FN, whereas GFP was enriched in the caudal-lateral FN, also known as the dorsal-lateral protuberance (DLP). Distinct regions devoid of fluorescence that correspond to the somata of individual CbN neurons were apparent in single confocal image plane (**Fig. 5A-C**).

**Figure 5.**
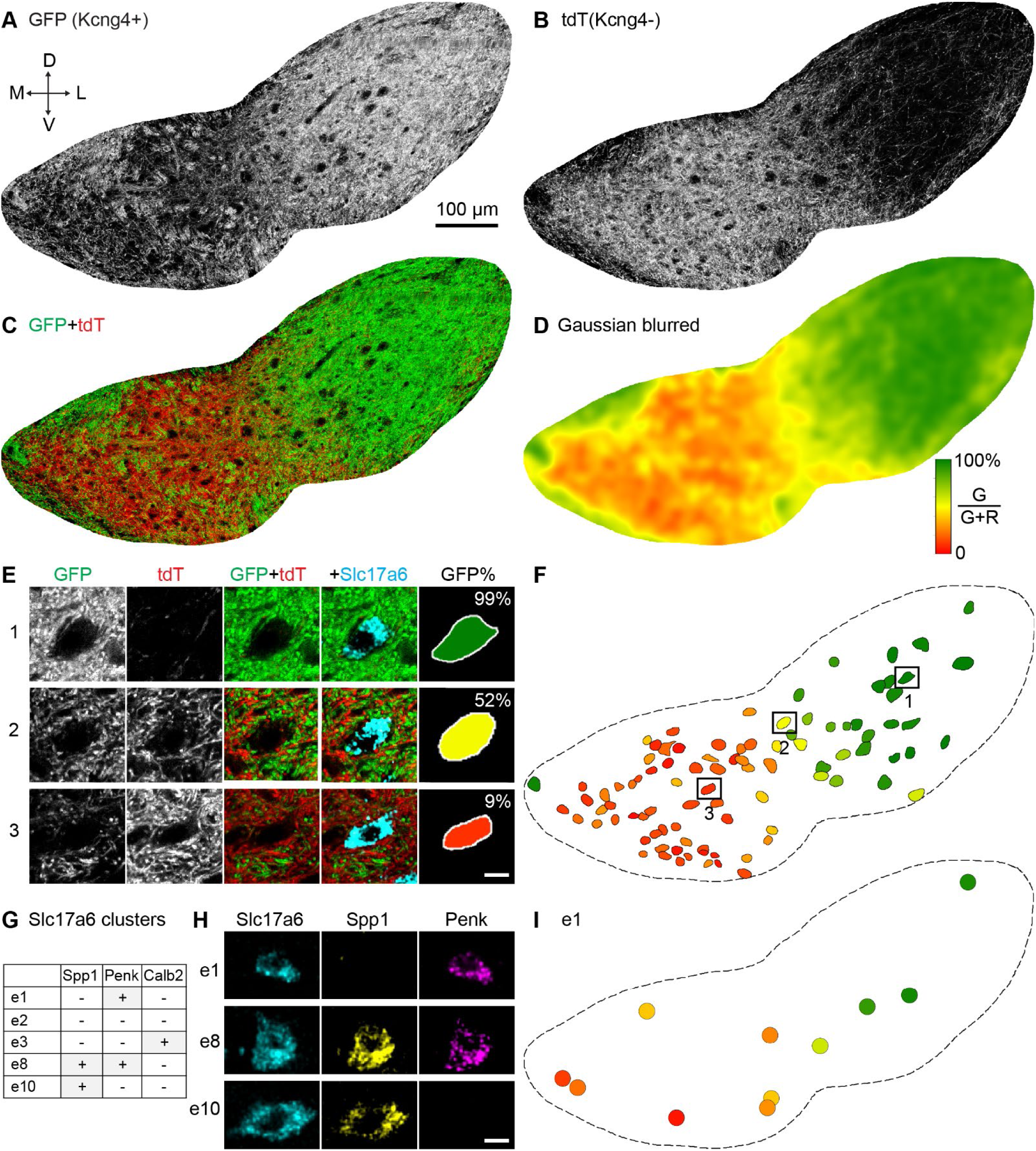
Quantifying PC projections to CbN neuron subtypes. The approach used to characterize PC subtype outputs is illustrated in a slice from a Kcng4^Cre^Pcp2^Flp^RC::FLTG mouse. **A.** Greyscale confocal image of GFP labelling of Kcng4+ PCs axons and boutons in the FN (single Z plane). **B.** As in **A** but for tdT labelling of Kcng4-PCs. **C.** Overlay of GFP (*green*, Kcng4+ PCs) and tdT labeling (*red*, Kcng4-PCs). **D.** Gaussian blur with the ratio of GFP to tdT labeling shown (based on a confocal stack). **E.** Labeling is shown for three Slc17a6+ cells used to determine the PC-subtype inputs the cells receive. **F.** Map of Slc17a6+ cells with each cell color coded based on the PC-subtype synapses on their cell bodies. **G.** Table of FN cell types and probes used to identify them. **H.** Examples of using ISH to characterize three types of FN neurons. **I.** Color-coded symbols showing G/(G+R) ratio for the e1 cells in **F**.

Substantial overlap between GFP and tdT signals was evident in the caudal-medial FN (**Fig. 5C**), which was readily visualized by applying a Gaussian blur to the GFP to tdT fluorescence ratio (**Fig. 5D**) to highlight regions dominated by GFP inputs (*green*), regions dominated by tdT inputs (*red*), and regions where both types of inputs are present (*yellow*). A similar analysis of a section from a Gpr176^Cre^Pcp2^Flp^RC::FLTG mouse showed that Gpr176+ and Gpr176-PC projections also exhibit considerable overlap in some regions of caudal FN (**Fig. S5**).

Interpretation of the images in **Fig. 5A-D** is complicated by the fact that both PC axons and boutons are fluorescently labeled, making it difficult to distinguish synaptic inputs from fibers of passage. We addressed this issue by quantifying the fluorescence of PC boutons made onto the somata of individual glutamatergic CbN neurons that were identified by *Slc17a6* (encoding VGLUT2) ISH, which are the principal projection neurons of the CbN (**Fig. 5E**). Some neurons received predominantly GFP inputs (**Fig. 5E**, *cell 1*), other exhibited a mixture of GFP and tdT inputs (*cell 2*), and others were primarily innervated by tdT inputs (*cell 3*). We used this approach to characterize all glutamatergic neurons within the section and generate a spatial map of *Slc17a6*+ somata color-coded by the ratio of different PC inputs onto their somata (**Fig. 5F**). There was a strong regional dependence of the cells targeted by different PC subtypes, and a mixture of inputs from both PC subtypes was still apparent for many cells, but the degree of convergence was reduced relative to **Fig. 5D**. To directly compare the estimates obtained from these two analyses, we compared the GFP input ratio obtained from somatic inputs as described in **Fig. 5E and 4F** to the ratio estimated from the Gaussian blur in **Fig. 5D**, which are expected to be more strongly influenced by fibers of passage and inputs onto neighboring cells. The somatic input quantification indicated a higher degree of segregation (**Fig. S6**), suggesting that it provides a more accurate estimate of PC subtype convergence onto individual CbN neurons.

The molecular identity of the CbN neurons is another vital issue in characterizing output pathways through the CbN. snRNA-seq studies have shown that there are fourteen types of excitatory neurons across all three CbN subnuclei ^20^. Here we focus on the FN that has five types of excitatory neurons (e1, e2, e3, e8, and e10) ^20^ that project to distinct extracerebellar regions implicated in diverse behaviors ^21^. Because these neuron types are often intermingled within the FN, it is necessary to determine the molecular identity of the targeted CbN neurons in addition to identifying the inputs from PC subtypes. We accomplished this by combining the characterization of PC subtype inputs (**Fig. 5E and 4F**) with ISH characterization of cell type based on markers previously identified in an snRNA-seq study ^20^ (**Fig. 5G**). This approach was illustrated for an e1 cell (*Slc17a6+, Spp1-, Penk+*), an e8 cell (*Slc17a6+, Spp1+, Penk+*), and an e10 cell (*Slc17a6+, Spp1+, Penk-*) (**Fig. 5H**). We applied this approach to the section in **Fig. 5F** and generated a map of all e1 cells, with each symbol color-coded to represent the ratio of Kcng4+ and Kcng4-inputs onto each cell (**Fig. 5I**). This map illustrates that e1 cells are present throughout the section and can be targeted primarily by either Kcng4+ or Kcng4-PC inputs depending on the location of the cell.

We applied this approach to characterize *Kcng4+* and *Kcng4-* projections onto the five molecularly defined excitatory neuron subtypes in the FN (**Fig. 6A**). Coronal sections spanning the FN were collected from a Kcng4^Cre^Pcp2^Flp^RC::FLTG mouse at the indicated planes (**Fig. 6A**, *upper left*). Gaussian-blurred images (as in **Fig. 5D**) provided an overview of the projection patterns of PC subtypes (**Fig. 6A**, *left*). We then quantified PC subtype inputs onto the somata of individual CbN neurons (as in **Fig. 5E and 4F**) and determined their molecular identities using ISH (as in **Fig. 5G and 4I**). This analysis generated spatial maps for five types of glutamatergic neurons across the selected FN sections, with each symbol color-coded to represent the ratio of Kcng4+ and Kcng4-inputs (**Fig. 6A**, *right*). Some glutamatergic neuron types exhibited strong regional distribution patterns – for example, e2 and e10 neurons were primarily localized to caudal FN, whereas e8 neurons were enriched in rostral FN – while others, such as e1 neurons, were more evenly distributed. Across all five neuron types, PC input patterns were largely determined by anatomical localizations: different types of neurons within a given region received inputs from PC subtypes that were consistent with the local PC projections patterns revealed by the Gaussian-blurred images, regardless of their molecular identities. The relative abundance of each glutamatergic FN neuron type is summarized in **Fig. 6C**. Because e2 neurons comprise a large fraction of glutamatergic FN neurons and occupy subregions implicated in distinct functional roles, we subdivided them into medial (cFN) and lateral groups (cDLP) in the two most caudal sections (**Fig. 6A**, Bregma -6.6 and -6.7, separated by dashed lines), and classified the remaining e2 neurons as rostral (**Fig. 6A**, Bregma -6.1, -6.3 and -6.4). These subdivisions were guided in part by a study characterizing projections made by different FN neuron types based on gene expression and distribution ^21^. We quantified the strengths of PC subtype output pathways by summing the fractions of Kcng4+ PC inputs across all neurons within each neuron subtype, as well as for Kcng4-inputs (**Fig. 6D**). This analysis indicated that Kcng4+ PCs account for 64% of the total PC outputs to glutamatergic FN neurons, whereas Kcng4-PCs account for the remaining 36%. Although both Kcng4- and Kcng4+ PCs provided inputs to all FN neuron types, they showed strong preferences: Kcng4+ PCs outputs are primarily mediated by e8, e10, and rostral e2 neurons, whereas Kcng4-PCs outputs are primarily mediated by cFN e2 neurons along with small contributions from other types of neurons.

**Figure 6.**
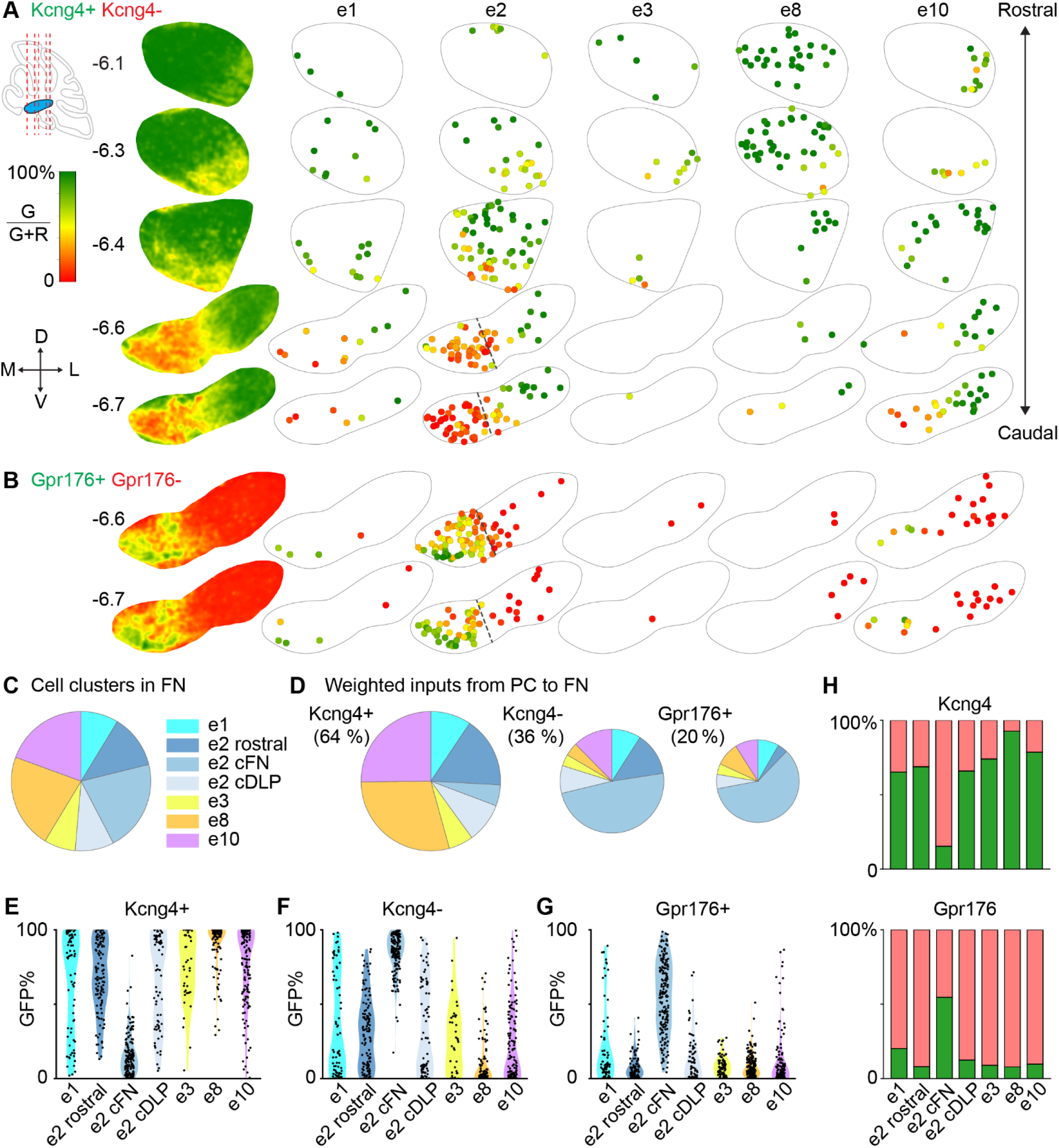
Projections of PC subtypes to different types of FN neurons. **A.** PC projections in the FN of Kcng4^Cre^Pcp2^FLP^RC::FLTG mice were characterized and types of Slc17a6+ cells were identified as in Fig. 5 for a series of slices at the indicated Bregma positions. The Gaussian blur images to the left are determined as in Fig. 5D and the spatial maps at the right are determined as in Fig. 5I, but for 5 types of excitatory cells. Dashed lines were used to subdivide e2 cells into e2 cFN and e2 cDLP in the -6.6 and -6.7 slices. **B.** Same as **A** but for Gpr176^Cre^Pcp2F^LP^RC::FLTG mice. There was very little GFP labeling in -6.1, -6.3 and -6.4 slices, so they are not shown. **C.** Summary of the fractional number of cells of each type. e2 cells were further subdivided based on region into e2 cFN, e2 cDLP (−6.6 and −6.7 slice) and e2 rostral (−6.0, -6.3 and -6.4 slices) cells. This plot is based on >1600 cells from 4 animals. **D.** The weighted PC inputs onto different subtypes is summarized. This was determined by adding the strengths of inputs onto each cell identified using ISH. This is based on 2 Kcng4 mice and 2 Gpr176 mice. **E.** Violin plot of the percentage of Kcng4+ inputs [green/(green +red)·100%] onto different types of FN neurons. **F.** Same as **E** but for Kcng4-PC inputs. **G.** Same as **E** but for Gpr176+ PC inputs. **H.** (*top*) Summary of the percentage of Kcng4+ PCs (green) and Kcng4-PCs (red) made onto different types of FN neurons. (*bottom*) Summary of the percentage of Gpr176+ PCs (green) and Gpr176-PCs (red) made onto different types of FN neurons.

We next applied the same approach to characterize Gpr176+ projections onto the glutamatergic FN neurons using Gpr176^Cre^Pcp2^Flp^RC::FLTG mice (**Fig. 6B**). Gpr176+ PCs provide prominent projections to the caudal FN (**Fig. 6B**, Bregma -6.6 and -6.7). The other sections are not shown because they are essentially devoid of Gpr176+ PC inputs. Gpr176+ PCs predominantly provide inputs to e2 neurons, which are the most abundant neuron type in the caudal FN, with additional but smaller contributions to e1 and e10 neurons. The weighted strengths of Gpr176+ PC inputs account for 20% of the total PC inputs onto glutamatergic FN neurons (**Fig. 6D**, *right*). This proportion likely exceeds the fraction of Gpr176+ PCs in the vermis (6.8%) due to the high density of glutamatergic neurons in caudal FN.

The composition of PC subtype inputs to individual identified FN neurons exhibited considerable cell-to-cell variability (**Fig. 6E-G**). Kcng4+ PCs provided most of the inputs to most classes of CbN neurons, with the exception of e2 cFN neurons, which mainly receive inputs from Kcng4-PCs (**Fig. 6E and 5F**). Many CbN neurons receive substantial convergent inputs from both Kcng4+ and Kcng4-PCs. The plot of Kcng4-inputs (**Fig. 6F**, the inverse of the Kcng4+ plot, **Fig. 6E**) is included for comparison with the Gpr176+ PC inputs (**Fig. 6G**). The very strong Kcng4-PC inputs (primarily Aldoc+ PCs) to e2 cFN neurons (**Fig. 6F**) and the weaker Gpr176+ PC inputs (Aldoc1) to e2 cFN neurons (**Fig. 6G**) indicate contributions from Aldoc+ subtypes other than Aldoc1. Gpr176+ PCs also provide strong inputs to a small subset of e1, e2 cDLP, and e10 neurons (**Fig. 6G**). These results indicate that the weighted inputs from PC subtypes to CbN neuron types (**Fig 5D**) reflect substantial contributions from individual CbN neurons with mixed PC subtype inputs. It is also informative to consider the percentage of PC subtype inputs to different types of FN neurons (**Fig. 6H**). Kcng4+ and Kcng4-PCs are strongly segregated for e2 cFN neurons, e8 and e10 neurons, but they converge to a substantial degree on other FN cell types. Gpr176+ PCs provide more than half of the inputs to e2 cFN neurons but contribute only a small fraction of the total inputs for other FN neuron types.

The identities of PC subtypes that form synapses onto distinct types of glutamatergic FN neurons provides insights into the functional roles of PC subtypes. Kcng4+ PCs extensively synapse onto e8, e10, and rostral e2 neurons, which have been shown to project to downstream regions implicated in motor and posture control ^21^. The other neuron types, especially those in cFN, receive inputs from Kcng4-PCs and Gpr176+ PCs and project to regions involved in vigilance. This suggests that PC subtypes defined by Kcng4 and Gpr176 could play differential roles in these behaviors.

### Silencing the outputs of Kcng4+ PCs selectively affects motor performance and motor learning

Chronically silencing all PC outputs leads to severe deficits in numerous motor and non-motor behaviors, including baseline motor performance, motor learning, sociability, and anxiety-like behaviors ^71,72^. Based on the distributions of PC subtypes defined by Kcng4 and Gpr176 in the cerebellar cortex and their distinct output pathways, we hypothesized that they regulate different behaviors. To test this hypothesis, we used the Cre/Flp-dependent RC::PFtox line to selectively express tetanus toxin in PC subtypes and silence their synaptic outputs. The RC::PFtox line has been extensively used to selectively silence the outputs of specific neuron subtypes for functional characterization ^73–76^. While the Pcp2^Flp^ line selectively targets PCs in the brain, it also labels a subset of retinal bipolar neurons ^77,78^. We therefore assessed whether the retina function and vision might be affected in our intersectional approach. In Kcng4^Cre^Pcp2^Flp^RC::FLTG mice, a small fraction of bipolar neurons is GFP+ (Pcp2+Kcng4+), and their synapses are expected to be silenced in Kcng4^Cre^Pcp2^Flp^RC::PFtox mice (**Fig. S7A**). Nonetheless, these mice showed normal performance in the optokinetic reflex (OKR) assay, in which the eyes generate an involuntary movement in response to a moving visual scene ^79,80^ (**Fig. S7C**). There are no Gpr176+Pcp2+ retinal cells (**Fig. S7B**), and OKR performance was normal in Gpr176^Cre^Pcp2^Flp^RC::PFtox mice (**Fig. S7D**). Together, these results suggest that the behavioral effects in these mice likely arise from silencing the synapses of targeted PC subtypes instead of retinal bipolar neurons.

We first characterized behaviors in Kcng4^Cre^Pcp2^Flp^RC::PFtox (Kcng4-tox) mice, in which synaptic outputs from Kcng4+ PCs were silenced. Kcng4+ PCs are mainly found in the anterior vermis and hemisphere of the cerebellar cortex (**Fig. 1** and **Fig. 2**), project to the rostral-dorsal part of the CbN (**Fig. 4**), and preferentially target e8, e10, and rostral e2 neurons in the FN (**Fig. 6**). This implicates Kcng4+ PCs in posture, balance, gait, and motor coordination. We therefore hypothesized that silencing Kcng4+ PC outputs might selectively impair motor function. Gait was analyzed by visualizing mice from below and two side views as they walked through a corridor and tracking multiple body parts (**Fig. 7A**) ^72,81,82^. Plots of the trajectories of these body parts across multiple gait cycles revealed that Kcng4-tox mice showed abnormal gaits compared to littermate controls (**Fig. 7B**). Kcng4-tox mice had increased fore-paw width and larger fluctuations for vertical rear, horizontal tail and vertical tail (**Fig. 7C**). Gait abnormalities in Kcng4-tox mice were similar to those present in mice where all PC outputs were silenced ^72^. Kcng4-tox mice also have impaired performance in the balance beam assay, with an increased traverse time and more paw slips (**Fig. 7D**). Performance on the rotarod assay was also impaired, although learning was still present in Kcng4-tox mice (**Fig. 7E**). These results show that Kcng4+ PCs are important for baseline motor performance as predicted by the location of Kcng4+ PCs and their outputs.

**Figure 7.**
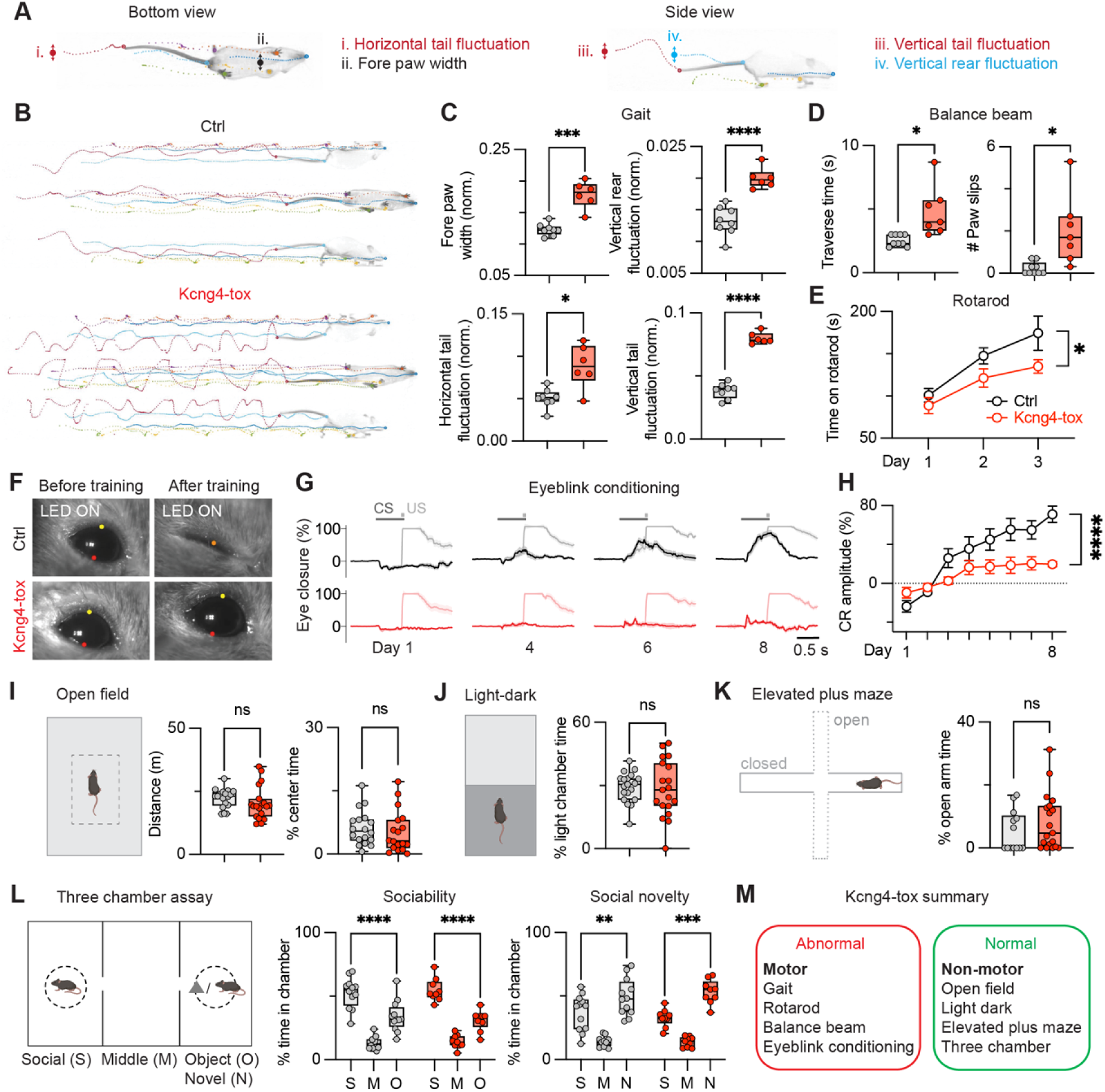
Kcng4-tox selectively affect motor performance and motor learning. **A.** Diagrams show the different body parts of mice extracted from a bottom view video and a side view video, and the parameters being measured in the gait analysis. **B.** Example gait tracking images of different body parts in ctrl and Kcng4-tox mice. **C.** Quantifications of fore paw width (norm.), vertical rear fluctuation (norm.), horizontal tail fluctuation (norm.), and vertical tail fluctuation (norm.) of ctrl (n=8) and Kcng4-tox (n=6) mice in the gait analysis. Only gait cycles with velocity 0.3-0.4 m/s are included in the analysis. **D.** Quantifications of traverse time (s) and number of paw slips of ctrl (n=9) and Kcng4-tox (n=7) mice in the balance beam assay. **E.** Quantifications of time on rotarod (s) across multiple days in ctrl (n=9) and Kcng4-tox (n=7) mice in the rotarod assay. **F.** Representative images showing the conditioned eyelid response to light (CS) before and after the training. Yellow and red dots represent upper and lower eyelid, while a single orange dot represent a fully closed eyelid. **G.** Averaged traces showing the eye closure (%) in the CS-US trials (light color trace) and CS-only trials (dark color trace) across multiple days in ctrl (n=11) and Kcng4-tox (n=10) mice. **H.** Quantifications of the conditioned response (CR) amplitude (%) in the CS-only trials across multiple days in ctrl (n=11) and Kcng4-tox (n=10) mice. **I.** *Left*, diagram showing the open field assay. *Right*, quantifications of total traveled distance (m) and center time (%) in ctrl (n=18) and Kcng4-tox (n=18) mice. **J.** *Left*, diagram showing the light-dark chamber assay. *Right*, quantification of light chamber time (%) in ctrl (n=19) and Kcng4-tox (n=19) mice. **K.** *Left*, diagram showing the elevated plus maze assay. *Right*, quantification of open arm time (%) in ctrl (n=13) and Kcng4-tox (n=19) mice. **L.** *Left*, diagram showing the three-chamber assay. *Right*, quantification of time in each chamber in sociability or social novelty assay in ctrl (n=12) and Kcng4-tox (n=9) mice. **M.** Summary of all the behavioral phenotypes in the Kcng4-tox mice.

We then looked at eyeblink conditioning, a widely studied behavioral assay for cerebellar-dependent motor learning ^83^. We hypothesized that conditioned eyeblink would be impaired in Kcng4-tox mice because this behavior involves paravermal lobules V and VI in the cerebellar cortex where Kcng4+ PCs predominate (**Fig. 1** and **Fig. 2**) and the rostral IP where Kcng4+ PC outputs are prominent (**Fig. 4**). After 8 days of training in which mice were presented with an LED signal that serves as a conditioning stimulus (CS), followed by an air puff to the left eye, which is the unconditioned stimulus (US) that evokes an eye blink, mice learned to respond to the LED with an anticipatory eyeblink (**Fig. 7F**, *top*; CS-only trials). Control mice gradually acquired the conditioned response (CR) in CS-only trials as early as learning day 3 and showed 80% eyelid closure in learning on day 8 (**Fig. 7F-H**; *black*), whereas Kcng4-tox mice failed to respond to the CS after 8 days of training (**Fig. 7F-H**; *red*). This establishes that Kcng4+ PCs are required for conditioned eyeblink learning as predicted.

We performed additional behavioral assays to gain insight into exploratory behaviors and anxiety-like behaviors that are known to be disrupted by silencing all PC outputs ^71^. We used open field assay to evaluate locomotion (total distance travelled) and used time spent in the center of the arena to evaluate exploratory behavior and anxiety-like behavior. Silencing all PC outputs decreased overall locomotion and distance travelled ^71^, which was not observed in Kcng4-tox mice despite their severe ataxia phenotypes (**Fig. 7I**). Kcng4-tox mice did not show altered time spent in the center of the arena, indicating that anxiety level and exploratory behavior were unaltered, as was the case when all PC outputs were silenced ^71^. We also assessed the innate aversion to bright light using a light-dark chamber assay and found that Kcng4-tox mice spent similar time in the light chamber as the control littermates (**Fig. 7J**). This contrasts with mice with all PC outputs silenced spent less time in the light chamber, which was consistent with elevated anxiety ^71^. Lastly, we found that Kcng4-tox mice spent similar time as the control littermates in the open arms of the elevated plus maze (**Fig. 7K**), again confirming that anxiety level and exploratory behaviors were not perturbed. Together, these results establish that silencing Kcng4+ PC outputs did not alter general locomotion, exploratory behaviors, and anxiety-like behaviors.

Many targeted manipulations of PCs lead to severe social deficits that are apparent in a three-chamber assay ^1,2,23,71,84,85^. In the sociability assay, a subject mouse is allowed to explore three interconnected chambers, with one chamber containing an unfamiliar mouse, and another containing an unfamiliar object (**Fig. 7L**, *left*). Mice normally prefer to explore the social chamber (S) rather than the object chamber (O) or middle chamber (M), but mice with all PC signaling silenced did not show a strong preference for the social chamber ^71^. Here, both Kcng4-tox and their control littermates showed strong preference for the social chamber (**Fig. 7L**, *middle*). We also assessed social novelty preference by keeping the mouse in the social chamber (it is now familiar) and replacing the object with a novel stranger mouse (N). Kcng4-tox mice and their control littermates both showed strong preference for the social novelty chamber (**Fig. 7L**, *right*). Therefore, silencing Kcng4+ PC outputs did not affect sociability or social novelty preference.

Together, these results establish that perturbing a PC subclass can selectively disrupt some behaviors but spare others. Suppressing Kcng4+ PCs selectively impaired numerous motor behaviors such as gait, balance beam, rotarod and conditioned eyeblink, but spared most cerebellar-dependent non-motor behaviors, including exploratory behaviors, anxiety-like behaviors, and social behaviors (**Fig. 7M**).

### Gpr176+ PCs are involved in anxiety-related and exploratory behaviors

We then characterized the same behaviors in Gpr176^Cre^Pcp2^Flp^RC::PFtox (Gpr176-tox) mice, in which synaptic outputs from Gpr176+ PCs were silenced. Gpr176+ PCs are most prominent in lobules VI, VII, and X of the vermis and in the PF (**Fig. 1 and 2**). Gpr176+ PCs project to caudal-medial FN, ventral-lateral IP, DN (**Fig. 4**), and more specifically, to the cFN e2 neurons (**Fig. 6**). These regions are implicated in diverse behaviors including vestibular processing and balance (lobule X) ^86^, cognitive function and emotional regulation (lobule VII) ^87^, saccades and oculomotor control (lobule VI/VII) ^88–90^, smooth pursuit eye movements (PF) ^91^ and vigilance (caudal-medial FN) ^21^. These findings suggest that disrupting Gpr176+ PC outputs might alter some specific behaviors but not baseline motor function.

Indeed, baseline motor functions were not perturbed in Gpr176-tox mice. Gait was normal (**Fig. 8A**) with unaltered fore paw width and tail fluctuations (**Fig. 8B**). Balance beam performance was also unaltered based on traverse time and the number of paw slips (**Fig. 8C**). The rotarod assay showed unaltered performance on day 1 and similar learning on subsequent days (**Fig. 8D**). In the eyeblink conditioning, both control and Gpr176-tox mice started to show learning of the conditioned eyeblink response starting from training day 3, and reached similar CR amplitude on training day 8 (**Fig. 8E-G**). These results establish that Gpr176+ PCs are not required for baseline motor performance and eyeblink conditioning. These findings are consistent with the observation that Gpr176+ PCs are not present in regions that contribute to motor function.

**Figure 8.**
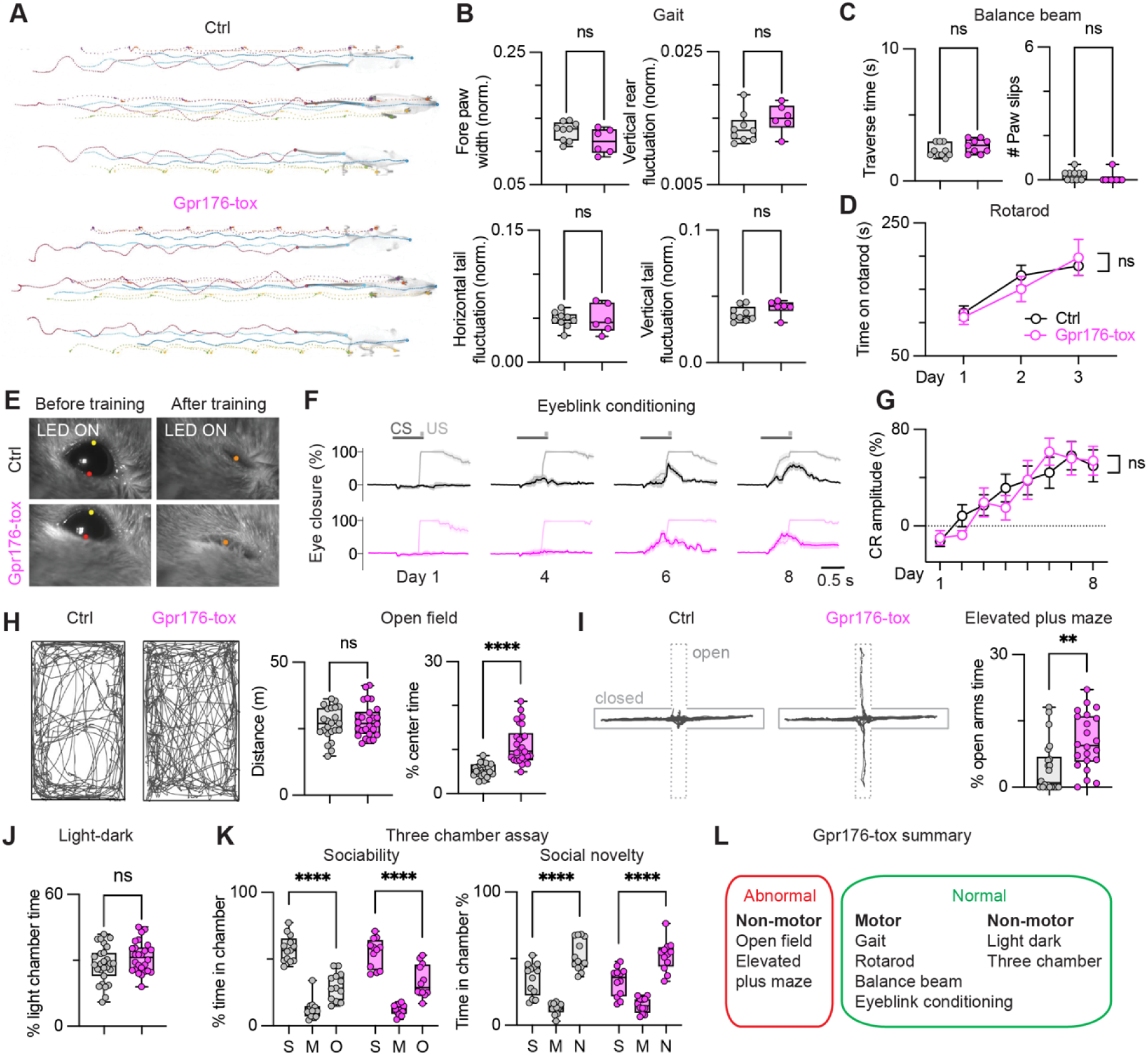
Gpr176-tox selectively affect exploratory behaviors. **A.** Example gait images in ctrl and Gpr176-tox mice. **B.** Quantifications of fore paw width (norm.), vertical rear fluctuation (norm.), horizontal tail fluctuation (norm.), and vertical tail fluctuation (norm.) in the gait analysis of ctrl (n=9) and Gpr176-tox (n=6) mice. Only gait cycles with velocity 0.3-0.4 m/s are included in the analysis. **C.** Quantifications of traverse time (s) and number of paw slips in the balance beam of ctrl (n=10) and Gpr176-tox (n=9) mice. **D.** Quantifications of time on rotarod (s) in the rotarod assay across multiple days in ctrl (n=10) and Gpr176-tox (n=9) mice. **E.** Representative images showing the eyelid response to light (CS) before and after the training. Yellow and red dots represent upper and lower eyelid, while a single orange dot represent a fully closed eyelid. **F.** Averaged traces showing the eye closure (%) in the CS-US trials (light color trace) and CS-only trials (dark color trace) across multiple days in ctrl (n=11) and Gpr176-tox (n=9) mice. **G.** Quantifications of the conditioned response (CR) amplitude (%) in the CS-only trials across multiple days in ctrl (n=11) and Gpr176-tox (n=9) mice. **H.** *Left*, representative tracking images of a ctrl and a Gpr176-tox mice in the open field assay. *Right*, Quantifications of total traveled distance (m) and center time (%) in ctrl (n=23) and Gpr176-tox (n=26) mice. **I.** *Left*, representative tracking images of a ctrl and a Gpr176-tox mice in the elevated plus maze assay. *Right*, Quantification of open arm time (%) in ctrl (n=22) and Gpr176-tox (n=22) mice. **J.** Quantification of light chamber time (%) in ctrl (n=27) and Gpr176-tox (n=26) mice in the light dark assay. **K.** Quantification of time in each chamber in sociability or social novelty assay in ctrl (n=13) and Gpr176-tox (n=12) mice in three-chamber assay. **L.** Summary of all the behavioral phenotypes in the Gpr176-tox mice.

Intriguingly, several non-motor tasks were selectively impaired in Gpr176-tox mice. In the open field assay, Gpr176-tox mice travelled the same overall distance but spent considerably more time exploring the center of the arena, as shown in representative tracking images and quantifications across many mice (**Fig. 8H**). In the elevated plus maze assay, Gpr176-tox mice spent much more time in the open arms than control mice (**Fig. 8I**). 55% (12/22) of control mice did not enter the open arms at all, compared to 14% (3/22) in Gpr176-tox mice. Interestingly, Gpr176-tox mice spent the same amount of time in the light chamber in the light-dark assay (**Fig. 8J**). Therefore, the Gpr176-tox mice showed increase exploratory behavior and reduced anxiety in the open field and elevated plus maze assay, while preseving the aversion to bright light in the light-dark assay. The increased exploratory behaviors of Gpr176-tox mice observed in the open field and elevated plus maze assays likely reflectes increased risk-taking and reward-seeking behaviors in a novel environment, in addition to a general decrease in anxiety. This behavioral effect is consistent with the presence of Gpr176+ PCs in lobule VI/VII, which is involved in emotion, and their projection to the cFN e2 neurons, which is implicated in vigilance ^21^. Lastly, in the three-chamber assay, social preference and novelty preference were normal in Gpr176-tox mice (**Fig. 8K**). This is not surprising, given that there are very few Gpr176+ PCs present in the primary region implicated in the regulation of social behaviors (Crus I) ^84,85^. Collectively, these findings establish that Gpr176+ PCs selectively regulate anxiety-like and exploratory behaviors, but are not required for general motor performance and many other cerebellar-dependent non-motor behaviors (**Fig. 8L**).

## Discussion

Intersectional PC subtype targeting allowed us to map their distributions within the cerebellar cortex and their outputs, and to quantify the synapses of PC subtypes on specific types of CbN neurons. By extending this approach to selectively silence PC subtype outputs we found that PC subtypes regulate specific cerebellar-dependent behaviors while sparing others. The approaches demonstrated here for Kcng4+ PCs and Gpr176+ PCs provide a versatile framework for investigating PC diversity, circuit specializations, and behavioral control.

### Selectively targeting PC subtypes

The key to our experimental strategy was to selectively target PC subtypes without affecting other types of neurons. This allowed us to perform behavioral experiments without invasive injections and accurately characterize PC outputs without interference from other brain regions. Generating a PC-specific Pcp2^Flp^ mouse was vital to our intersectional approach (**Fig. 1A**, **Fig. S1A**). Combining Pcp2^Flp^ mice with subtype-specific Cre driver lines allowed us to exclusively target PC subtypes in the brain. In Kcng4^Cre^Pcp2^FLP^RC::FLTG mice, Kcng4+ and Kcng4-PCs were widely distributed across all cerebellar lobules in alternating parasagittal stripes (**Fig. 1 and 2**). Kcng4+ PCs predominate in the anterior vermis and hemisphere, whereas Kcng4-PCs are enriched in the posterior vermis, Crus I and PF. Within the vermis and hemispheres, Kcng4+ PCs largely correspond to Aldoc-PCs. Nonetheless, there are minor differences between PCs labeled with these two markers, and other *Aldoc+* markers (*Plcb3* and *Slc1a6*) ^16,50–53^. These differences indicate that while subdividing PCs into two classes is a useful approximation, there is additional heterogeneity within each class, particularly in the transition zones.

We also generated a Gpr176^Cre^ line to target Aldoc1 PCs, which have a distinctive molecular profile that differs in interesting ways from other Aldoc+ subtypes ^15^, as illustrated by the differential expression of several genes implicated in synaptic plasticity, including *Plcb1*, *Plcb4*, and *Prkca*, that are expressed at extremely low levels in Aldoc1 PCs, intermediate levels in other Aldoc+ PCs, and very high levels in Aldoc-PCs (**Fig. 1B**). We found that Gpr176+ PCs are organized into discrete stripes or patches in specific cerebellar regions (**Fig. 1 and 2**). Gpr176+ PCs are abundant in lobule VI (14.7% of PCs) that contribute to oculomotor control and fear memory consolidation ^92^, lobule VII (18.1%) that regulates anxiety and exploratory behavior ^87^, lobule X (31.9%) that is involved in vestibular processing ^93,94^, and the PF (51.3%) and F (19.1%) that integrates vestibular and visual inputs ^91^. Their distinctive distribution pattern and low expression of key plasticity-related genes suggest that Gpr176+ PCs contribute to specialized information processing in multiple regions to regulate diverse behaviors.

The approach we developed can be readily extended to target additional PC subtypes. A single cross of an appropriate Cre driver mouse with a Pcp2^Flp^RC::FLTG mouse generates progeny that enable immediate visualization of the distribution of a PC subtype (GFP) and all other PCs (TdT). The unfolding algorithm we developed generates a 2D map of the entire cerebellar cortex from serial sections (**Fig. 2**; **Fig. S3**), which can be used to evaluate Cre line specificity and determine PC subtype distribution, thereby offering important clues about their potential behavioral functions.

### PC subtype properties

Our finding that the firing rates of Gpr176+ (Aldoc1) PCs in lobule X are lower than other Aldoc+ PCs (**Fig. 3**) establishes that the properties of PCs cannot be reduced to a simple Aldoc+ versus Aldoc-dichotomy; the particular molecularly defined PC subtype matters. Beyond firing rates, Aldoc+ and Aldoc-PCs differ in many additional ways. For example, EAAT4 levels are higher in Aldoc+ PCs, which alters the time course and amplitude of glutamate signaling, limits mGluR1 activation of TRP receptors, limits endocannabinoid release from PCs, and reduces the magnitude of CF-MLI spillover synapses ^32,57,58^. Given the differential expression of multiple genes in the mGluR1 signaling pathway (*Plcb1, Plcb3, Plcb4*, and *Prkca*; **Fig. 1B**) between Aldoc1 and other Aldoc+ PCs, it is also likely that additional aspects of the firing and synaptic properties of Aldoc1 PCs also differ from other Aldoc+ PC subtypes.

### PC subtype outputs

The intersectional labeling strategy has numerous advantages for characterizing PC subtype outputs. Restricting expression to PCs avoids labeling extracerebellar neurons that project to the CbN or brainstem, in contrast to immunohistochemical or non-specific transgenic approaches. The complementary two-color labeling of distinct PC populations enables precise assessment of both overlap and segregations of PC projections in the downstream regions. Although Kcng4+ and Kcng4-PCs predominantly innervate different CbN regions, there is substantial overlap in the transition zones (**Fig. 5A-D**). Overlap of PC subtype projections was also evident in Gpr176^Cre^Pcp2^Flp^RC::FLTG mice (**Fig. S5**). This suggests that PC subtypes defined by Kcng4 and Gpr176 provide convergent inputs to specific regions of the FN.

Bright dual-fluorescence labelling also facilitated the use of confocal microscopy to quantify PC subtype inputs onto the somata of individual CbN neurons (**Fig. 5E**). This approach established that different PC subtypes provide inputs to single CbN neurons, which does not reflect fluorescence from nearby synapses or fibers of passage. The PC synapses onto the somata of CbN neurons are likely highly effective at suppressing firing of CbN neurons, but PC synapses onto dendrites are excluded from the analysis. Dendritic inputs could further increase the diversity of PC subtype inputs to individual CbN neurons.

By combining PC subtype input characterization with ISH, we could estimate the relative weights of different PC subtype synapses onto molecularly and regionally defined types of FN neurons (**Fig. 5G-I**, **Fig. 6**). This allowed us to quantitatively estimate the strengths of different output pathways for each PC subtype (**Fig. 6**). This approach can be readily extended to the interposed and dentate nuclei using probes tailored to those regions. It can also be combined with viral injections to characterize PC subtype projections from localized regions of the cerebellar cortex.

The properties of PC subtype-specific output pathways provide insight into their functional and behavioral roles. Kcng4+ PCs provide the vast majority of PC synapses onto e8 (93 %) and e10 (79 %) neurons, whereas Kcng4-PCs (85 %) and Gpr176+ PCs (55%) provide most of the inputs to e2 (cFN) neurons (**Fig. 6**). Both Kcng4+ and Kcng4-PCs contribute substantially to synapses onto e1, e2 (rostral), e2 (cDLP), and e3 neurons, suggesting that they jointly regulate behaviors controlled by these CbN neurons. It appears that Kcng4+ PCs and Kcng4-PCs are each specialized to selectively regulate some behaviors (**Fig. 7**), but they may also work together to regulate other behaviors. Our results can be directly compared with a recent anatomical study that characterized FN neuron subtypes in detail in terms of their downstream projections and behavioral implications, although our cell type classification was based on a snRNA-seq study that differs from their labeling convention ^20,21^. For the primary FN targets of Kcng4+ PCs, e8 neurons project to spinal cord, lateral and inferior vestibular nuclei, and nucleus reticularis gigantocellularis that are implicated in posturomotor control; e10 neurons project to brainstem regions associated with oromotor control, including the intermediate reticular nucleus and facial motor nucleus ^21^. For the primary FN targets of Kcng4-PCs and Gpr176+ PCs, e2 (cFN) neurons project to the centrolateral, medialdorsal, and ventromedial thalamus that are involved in cognition, arousal, and motor planning, as well as brainstem nuclei implicated in vigilance and neuromodulation, including the supramammillary region, substantia nigra pars compacta, and nucleus incertus ^21^. For FN neuron types that are targeted by both Kcng4+ and Kcng4-PCs, e2 (rostral) neurons are associated with motor function, e2 (cDLP) with threat, salience, and behavioral flexibility, and e3 neurons with locomotion-respiration coupling and posture-autonomic control ^21^. It is anticipated that future studies that precisely map projections from molecularly distinct neurons in all CbN nuclei will further clarify how PC subtype-specific output pathways shape behavior.

Our findings provide additional insight into multimodal integration in the cerebellum and the extent to which individual CbN neurons integrate inputs from PCs in different regions. Previous studies have reached divergent conclusions regarding this issue. Images of Aldoc+ PC outputs alone ^37,41^ concluded that Aldoc+ and Aldoc-PC projections to the CbN are strongly segregated. The simplest explanation for the extensive overlap of PC subtype inputs to CbN neurons seen in our studies, versus the reported segregation of Aldoc+ and Aldoc-PC inputs, is that they reflect differences between PCs labeled by *Kcng4* or *Aldoc*. It is also possible that imaging a single type of PC may underestimate the extent of overlap that becomes apparent when projections of other PC subtypes are visualized simultaneously, as we found for Kcng4+ and Kcng4-PCs (**Fig. 5** and **Fig. 6**). Single-PC labeling has shown that PCs within the same rostrocaudal Aldoc+ or Aldoc-stripe project to a small region in the CbN, whereas PCs from different mediolateral stripes innervate segregated CbN areas ^38^. PCs in transition zones between Aldoc+ and Aldoc-regions have not been studied extensively, and they might make disproportionately large contributions to overlapping projections. A recent study found that the convergence of PCs on CbN neurons is greater than had been appreciated ^95^. Optogenetic activation of PCs in different cerebellar lobules found that many CbN neurons receive inputs from multiple lobules and in some cases as many as 5 different lobules that span across Aldoc+ and Aldoc-zones. It will be important to extend these studies by determining the molecular identities of PCs in different lobules that converge onto the same CbN neuron to provide further insight into multimodal cerebellar integration.

### Comparison of the behavioral effects of silencing all PCs, Kcng4+ PCs, and Gpr176+ PCs

Our findings provide direct evidence that specialized PC subtypes regulate distinct cerebellar-dependent behaviors. Silencing all PC synapses in PC-TKO mice produced broad deficits in baseline motor performance, motor learning, exploratory behaviors, anxiety-like behaviors, and social behaviors ^71,72^, whereas silencing synapses in PC subtypes has more selective effects (**Table 1**). In Kcng4-tox mice, motor performance and motor learning were impaired, but cerebellar-dependent non-motor behaviors were unaffected. This pattern is consistent with the distribution of Kcng4+ PCs and their projections to the rostral-dorsal CbN and lateral VN. This suggests that the social and emotional behaviors spared in Kcng4-tox mice are likely mediated by Kcng4-PCs, which can be examined by silencing the outputs of Kcng4-PCs.

**Table 1.** Behavior summary for PC-TKO, Kcng4-tox and Gpr176-tox mice. In PC-TKO mice, CaV2.1, CaV.2.2 and CaV2.3 calcium channels have been selectively eliminated from PCs to selectively silence all PC synapses. The summary provided for PC TKO mice is based on Lee et al, 2023 and Lee et al., 2025.

| Behavioral assay | PC-TKO | Kcng4-tox | Gpr176-tox |
| --- | --- | --- | --- |
| Gait | Abnormal | Abnormal (Fig. 7B-C) | Normal (Fig. 8A-B) |
| Balance beam | Abnormal | Abnormal (Fig. 7D) | Normal (Fig. 8C) |
| Rotarod | Abnormal | Abnormal (Fig. 7E) | Normal (Fig. 8D) |
| Eyeblink conditioning | Abnormal | Abnormal (Fig. 7F-H) | Normal (Fig. 8E-G) |
| Open field, distance | Abnormal | Normal (Fig. 7I) | Normal (Fig. 8H) |
| Open field, center time | Normal | Normal (Fig. 7I) | Abnormal (Fig. 8H) |
| Light-dark chamber | Abnormal | Normal (Fig. 7J) | Normal (Fig. 8J) |
| Elevated plus maze | -- | Normal (Fig. 7K) | Abnormal (Fig. 8I) |
| Three-chamber assay | Abnormal | Normal (Fig. 7L) | Normal (Fig. 8K) |

In sharp contrast, Gpr176-tox mice exhibited normal motor and social behaviors but showed alterations in anxiety and exploratory behavior as assessed with three different assays. In the light-dark assay, silencing all PC synapses in PC-TKO mice decreased time spent in the brightly illuminated chamber, but Gpr176-tox mice were unaffected, indicating that the increased aversion to bright light in PC-TKO mice is driven by non-Gpr176 PCs. In the open field assay, center time was unchanged in PC-TKO mice but increased in Gpr176-tox mice, indicating an increase in exploratory behavior and reduced anxiety. This surprising result indicates that in some circumstances silencing the synapses of a single PC subtype can have a much larger behavioral effect than silencing the synapses of all PCs, and that non-Gpr176 PCs can mask the behavioral contributions of Gpr176+ PCs. In the elevated plus maze assay,

Gpr176-tox mice spent more time in the open arms, consistent with increased exploration and reduced anxiety. Our findings indicate that distinct PC subtypes differentially influence specific aspects of anxiety and exploratory behavior that are regulated by the cerebellum ^87^. The anxiety-related phenotypes in Gpr176-tox mice are similar to the effects of midline lesions of the entire vermis of 10 day old rats, which increased exploration and time spent in the open arms of elevated plus maze in adulthood ^96^, and to the effects of chronic elevation of PC firing in lobule VII, which decreased open arm time in the elevated plus maze ^55^. It is likely that Gpr176+ PCs in lobule VII mediate many of the effects seen in previous studies. The contrasting phenotypes of PC-TKO and Gpr176-tox mice suggest that multiple cerebellar pathways act in parallel to regulate these behaviors, potentially exerting synergistic or opposing effects depending on the specific circuit engaged. Further studies are required to dissect the differential contributions of Gpr176+ and other PC subtypes to these behaviors.

### Future studies of PC subtypes

Intersectional targeting of PCs is needed to understand how different PC subtypes are specialized to support diverse functions. Slice experiments indicate that synaptic properties and intrinsic firing of Aldoc+ and Aldoc-PCs differ in many ways ^30,32,35,97^, but nothing is known about specific PC subtypes. Differential gene expression suggests substantial diversity among the nine PC subtypes ^15^. Additional recordings from fluorescently labelled PC subtypes in brain slices will allow characterization of synaptic properties, local circuitry, long-term synaptic plasticity, and long-term plasticity of intrinsic excitability. *In vivo* recordings from optotagged PC subtypes can be used to assess the firing of different PC subtypes during behavior. In addition, local viral injections can be used to examine how particular PC subtypes within a defined cerebellar region contribute to specific behaviors. The behavioral phenotypes observed in Kcng4-tox and Gpr176-tox mice reflect chronic silencing of PC synapses that approximates PC loss in neurological disorders affecting PC viability. Combining intersectional targeting of PC subtypes with optogenetic and chemogenetic approaches will allow direct comparison of acute versus chronic perturbations, providing insights into the plasticity and adaptability of cerebellar circuits to prolonged perturbations.

## Methods

### Mice

Animal procedures were carried out in accordance with the NIH and Institutional Animal Care and Use Committee (IACUC) guideline, and protocols approved by the Harvard Medical Area Standing Committee on Animals. Both male and female mice were used in the experiments. Mice were breed on a mix background (C57BL/6 and FVB/NJ). Mice were housed under standard conditions in groups of 2-5 animals on a 12 h light-dark cycle at an ambient temperature of 22°C and ambient relative humidity of 50%, with food and water available *ad libitum*.

Pcp2^Flp^ and Gpr176^Cre^ knock in mice were generated in this study using Easi-CRISPR ^98^. In brief, long single-stranded DNAs (ssDNAs, GeneScript) doners containing either a Flp recombinase or a Cre recombinase flanked by homologous arms targeting the stop codon of the gene of interest (**Fig. S1A and S1B**) were injected together with Cas9 protein (PNA Bio) and single-guide RNA (sgRNA, Synthego) into FVB/NJ embryos by the Transgenic Core at BIDMC. Other mouse lines used in this study include Kcng4^Cre^ (^54^; RRID:IMSR_JAX:029414), Ai14 (RRID:IMSR_JAX:007914), RC::FLTG (^49^; RRID:IMSR_JAX:026932), RC::PFtox (^73^ MGI:4412286). Kcng4^Cre^ heterozygous and Gpr176^Cre^ heterozygous were crossed with Pcp2^Flp^RC::FLTG mice to generate Kcng4^Cre^Pcp2^Flp^RC::FLTG and Gpr176^Cre^Pcp2^Flp^RC::FLTG (triple-heterozygous) mice. Kcng4^Cre^ heterozygous and Gpr176^Cre^ heterozygous were crossed with Pcp2^Flp^RC::PFtox mice to generate Kcng4^Cre^Pcp2^Flp^RC::PFtox and Gpr176^Cre^Pcp2^Flp^RC::PFtox (triple-heterozygous) mice. Pcp2^Flp^RC::PFtox littermates were used as controls in behavioral experiments.

## Tissue processing

### Vibratome slicing and imaging

8–24-week-old mice were used in the study. Mice were anesthetized with an intraperitoneal injection of a Ketamine (100 mg/kg) and xylazine (10 mg/kg) mixture and perfused with ice cold phosphate buffered saline (PBS, pH = 7.4, Sigma Cat# P-3813), followed by a solution containing 4% paraformaldehyde in PBS (Thermofisher Cat# J19943.K2). Brains were removed and postfixed in the same solution at 4°C overnight. Brains were sliced (50 μm thick) in PBS using a vibratome (VT1000S, Leica). Slices were then mounted on slides (Superfrost Plus, VWR, Cat# 48311-703) and covered with mounting medium (Fluoromount-G with DAPI, Invitrogen Cat# 00-4959-52) and a glass coverslip. Images were acquired by Olympus VS200 slide scanners with a 10x lens.

### Cryostat slicing and HCR RNA FISH

20–24-week-old mice were anesthetized and perfused as described above. Brains were harvested and post-fixed in 4% PFA in PBS solution overnight, followed by cryoprotection in 15% sucrose and then 30% sucrose in PBS. Tissues were embedded in Tissue-Tek O.C.T. compound (VWR Cat# 25608-930), frozen on dry ice, and sectioned with a cryostat (CM3050S, Leica). Coronal sections (30 μm) containing the fastigial nuclei were collected onto slides (Superfrost Plus, VWR, Cat# 48311-703) and stored in -80°C until further processing. For the cryosection of retina, eyes were harvested and post-fixed in 4% PFA in PBS solution for 30 mins. The cornea was cut and the lens was removed before the routine cryoprotection procedure.

For HCR RNA FISH, slides were first fixed in 4% PFA in PBS for 15 minutes at 4°C, followed by sequential washes in 50%, 75%, and twice in 100% ethanol (5 minutes each) at room temperature, and then a 5-minute wash in PBS. Slides were incubated in Probe Hybridization Buffer (Molecular Instruments) at 37°C for 10 minutes followed by overnight hybridization at 37°C with probes at 1:250 dilution.

Following hybridization, slides were washed at 37°C in graded mixtures of Probe Wash Buffer (PWB, Molecular Instruments) and 5X SSCT (75% PWB/25% 5X SSCT, 50% PWB/50% 5X SSCT, 25% PWB/75% 5X SSCT, and 100% 5X SSCT, 15 minutes each), followed by an additional 5-minute wash in 100% 5X SSCT at room temperature.

Slides were then incubated in Amplification Buffer (Molecular Instruments) for 30 minutes at room temperature. During this time, hairpin amplifiers were snap-cooled (95 °C for 90 seconds and cooled to 20 °C). The slides were then incubated with the amplifiers diluted 1:50 in Amplification Buffer at room temperature overnight. The following day, the slides were washed in 5X SSCT twice for 30 minutes followed by a final 5-minute wash in 5X SSCT, and then mounted with Fluoromount-G without DAPI (Invitrogen, Cat# 00-4958-02).

## Image processing

### 3D reconstructions of the cerebellum

To generate 3D reconstructions of the cerebellum, images of serial 50-μm coronal sections were first cleaned and aligned using the BrainJ package in Fiji/ImageJ (https://github.com/lahammond/BrainJ, ^99^). The 3D volume of the aligned images was then visualized with Imaris. For 3D reconstructions of the CbN and brainstem, the same 3D volume was used, but the area outside of CbN or brainstem were masked out to show the PC axons signals inside CbN or brainstem. The 3D reconstructions were aligned to the reference Allen Brain Atlas using the ABBA package in Fiji/ImageJ (https://github.com/BIOP/ijp-imagetoatlas, ^100^).

### 2D maps of the cerebellar cortex

2D maps of the cerebellar cortex were generated using a custom Python pipeline. Serial 50-μm sagittal sections from one hemisphere of the cerebellum were analyzed. The DAPI channel was used to segment Purkinje cell layer (PCL) contours and identify valley points corresponding to lobule boundaries (**Fig. S3A-C**), while the GFP and tdT channels were used to segment PC bodies (**Fig. S3D and S3E**). These steps produced a matrix representation of the PCL containing information on length, lobular organization, and cell labeling patterns (**Fig. S3F**), which was visualized as a linearized (unfolded) PCL (**Fig. S3G**). This procedure was repeated for all sagittal sections, generating a series of matrices that were converted into aligned linear representations and centered at their midpoints to construct a 2D map of one cerebellar hemisphere. In this map, line width corresponds to section thickness (50 μm), and line length reflects the true length of the unfolded PCL, determined by the number of matrix points. The hemispheric map was then mirrored across the midline to produce a full 2D cerebellar map that preserves the original aspect ratio (**Fig. S3H**). The flocculus and paraflocculus were processed separately and manually positioned onto the final map at their corresponding anatomical locations.

### Confocal image acquisition and processing

Images were acquired using a Leica Stellaris 5 confocal microscope with a 63X objective. FN images were collected in photon-counting mode 16-bit depth with 4-8 frame accumulations to improve signal intensity. Four z-planes spaced 4 µm apart were acquired and exported as TIFF files.

The Slc17a6 ISH channel was segmented using Cellpose-SAM ^101^. followed by manual curation to define the regions of interest (ROIs) corresponding to individual excitatory neurons of FN.

GFP and tdT channels were background-corrected using signal intensities measured within ROIs, where neuronal somata are located and PC axons are largely absent. The signal intensities of GFP and tdT channels were then normalized to a range between 0 and the average intensity of top 5% pixels to enable cross-channel comparison. In rostral FN slices in Gpr176^Cre^Pcp2^Flp^RC::FLTG mice, where there were almost no Gpr176+ axons hence with little GFP signals, normalization was performed using intensities values from caudal sections of the same animal.

### Gaussian blur of the CbN projections

The GFP and tdT channels from individual z planes were processed using a Gaussian filter from SciPy. The filtering parameters are sigma = 20 pixels (translates to ∼7.2 µm), order = 0, mode = “reflect”, truncate = 4.0. Pixel-wise GFP percentage was calculated as GFP / (GFP + tdT) and rendered using a red-yellow-green color map for visualization.

### Quantification of PC inputs to individual CbN neurons

ROIs corresponding to excitatory FN neurons were expanded by 3 µm to capture surrounding axon terminals. Pixels with both GFP and tdT signal intensities under 1000 were excluded as background. Remaining pixels were classified as “green” or “red” depending on whether GFP or tdT signal predominated, and proportion of green pixels was calculated for each ROI.

### *In vitro* recording

Acute parasagittal slices (200-µm thick) were prepared from 3-4-month-old Gpr176^Cre^Pcp2^Flp^RC::FLTG mice. Mice were anesthetized with an intraperitoneal injection of ketamine (100 mg/kg) and xylazine (10 mg/kg) and perfused with an ice-cold solution containing (in mM): 110 choline chloride, 7 MgCl_2_, 2.5 KCl, 1.25 NaH_2_PO_4_, 0.5 CaCl_2_, 25 glucose, 11.6 sodium ascorbate, 3.1 sodium pyruvate and 25 NaHCO_3_, equilibrated with 95% O_2_ and 5% CO_2_. Slices were cut in the same solution and transferred to artificial cerebrospinal fluid (ACSF) containing (in mM): 125 NaCl, 26 NaHCO_3_, 1.25 NaH_2_PO_4_, 2.5 KCl, 1 MgCl_2_, 1.5 CaCl_2_ and 25 glucose, equilibrated with 95% O_2_ and 5% CO_2_. Following incubation at 34 ℃ for 30 min, the slices were kept up to 4 h at room temperature until recording.

Recordings were made from PCs in the vermal lobule X at 33.3 ± 0.1 ℃. Visually guided on-cell recordings were obtained with patch pipettes of 1-3 MΩ pulled from borosilicate capillary glass (BF150-86-10, Sutter Instrument). Gpr176 expression was confirmed by observation of GFP expression in the PCs. Spontaneous action potentials were recorded in loose-patch configuration with glass pipettes filled with ACSF for 3 min. Electrophysiology data were acquired using a Multiclamp 700B amplifier (Axon Instruments), digitized at 20 kHz and filtered at 4 kHz using Igor Pro (Wavemetrics) running mafPC (courtesy of M. A. Xu-Friedman). Acquisition and analysis of slice electrophysiological data were performed using custom code written in Igor Pro and MATLAB. To block glutamatergic and GABAergic synaptic currents, the following receptor antagonists were added to the ACSF solution (in µM): 2 (*R,S*)-CPP, 5 NBQX and 10 SR95531 (gabazine). All drugs were purchased from Abcam and Tocris.

Spikes from slice recordings were detected and analyzed offline in MATLAB. Firing rates were calculated as the number of all spike events divided by recording time. Coefficient of variation (CV) of inter-spike intervals (ISIs) were calculated as the standard deviation of the ISIs divided by their mean. PCs with a CV less than 0.2 were defined to have regular firing patterns, while PCs with a CV more than 0.2 were defined to show irregular firing patterns. Local CV (CV2) was calculated as CV2_i_= 2|ISI_i+1_-ISI_i_| / (ISI_i+1_+ISI_i_), and the overall CV2 value was obtained by averaging CV2_i_ across all consecutive ISI pairs. For visualization and comparison of spike waveforms, all spike events from each cell were averaged to obtain a mean spike waveform for each cell. These mean spike waveforms were ranked by the spike amplitude in each group, and the top six cells with the largest negative deflections were selected. Their waveforms were normalized so that the most negative value was -1, and averaged within each group.

### Behavior

8–16-week-old mice were used in the study. All behavioral testing was performed in both male and female mice with genotypes blinded to the experimenter. Before behavioral testing, mice were handled by experimenter for two days (15 minutes per day). All behavioral testing was performed in the dark cycle, with ambient light and white noise. On the test day, mice were allowed to acclimate in the behavioral room for at least 30 minutes before starting the experiments.

### Gait analysis

Gait analyses were performed as previously described ^82^. Mouse locomotion was recorded using a high-speed camera (Bonito CL-400B/C, Allied Vision; 2320 × 700 pixels, 200 frames per second) positioned beneath a transparent glass floor as mice walked through a linear corridor illuminated with infrared light. Two mirrors placed along the sides enabled simultaneous views from below and from both lateral perspectives. To encourage traversal, nesting material and food were placed at one end of the corridor (habituation side), while aversive stimuli (light and gas) were applied at the opposite end (aversive side). Each trial ended when the mouse reached the habituation side, and the video was saved. Eight trials per mouse were conducted daily over five consecutive days.

Videos were analyzed using a convolutional neural network to track body parts (nose, left and right forepaws, left and right hind paws, rear end, and tail tip), as previously described. A hidden Markov model was then applied to segment the trajectories into individual gait cycles. All segmented cycles were manually reviewed by an experimenter blinded to genotype. Cycles were excluded if body parts were incorrectly labeled or if the mouse paused or moved backward during the cycle. We quantified the following gait parameters: cadence (cycles per second); stride length (maximum distance traveled by a forepaw within a single cycle); fore- and hind paw width (lateral distance between the forepaws or hind paws); and velocity (average speed of the mouse’s body center, defined along the corridor axis, within a single gait cycle). We also measured tail and rear height (absolute height of the tail tip and rear end above the floor) and tail and rear fluctuation (standard deviation of horizontal or vertical movement). Stride length, paw width, and tail and rear height and fluctuation were normalized to body length. For statistical analysis, gait parameters were averaged across cycles for each mouse. All analyses were performed using custom MATLAB code.

### Balance beam

Mice traversed a narrow beam (60 cm long, 1 cm wide) elevated 30 cm above the floor. To encourage movement across the beam, a dark box containing nesting material and food was placed at one end, while a bright aversive light (>600 lux) was positioned at the starting end. Mice were acclimated in the goal box for 2 minutes before training.

During training, mice completed four trials, starting at increasing distances from the goal box (10, 20, 30, and 40 cm), to learn to traverse the beam. During testing, left- and right-side views were recorded using two cameras (ELP, Ailipu Technology Co.; 30 Hz, 640 × 480 pixels) with iSpy open-source software. Mice were allowed to rest for 30 seconds between trials and were tested until they completed three uninterrupted trials (no pauses >2 s). For analysis, the number of slips of the left and right hind paws and the traversal time were manually scored by an experimenter blinded to genotype, and values were averaged across the three trials.

### Rotarod

Mice were tested on an accelerating rotarod (Rotamex-5, Columbus Instruments) using Rotamex-5 software. The rod speed increased from 4 to 40 rpm in 1 rpm increments every 8 seconds. Falls were detected automatically by infrared photocells positioned above the rod, and latency to fall was recorded. A fall was defined as either the mouse dropping off the rod or passively rotating (clinging and looping) without running. Mice completed three trials per day over three consecutive days, with a 1-minute rest period between trials. For each day, the latency to fall was averaged across the three trials.

### Eyeblink conditioning

Eyeblink conditioning was performed as previously described ^71^. Five days prior to the experiment, a head bracket was surgically implanted on the skull of each mouse. Animals then underwent one day of habituation to acclimate them to the apparatus. During habituation, mice were head-fixed for 20 minutes on a motorized treadmill (6-inch diameter) rotating at 2.5 cm/s. During training, a 550 ms white LED flash delivered to the left eye served as the conditioned stimulus (CS) and co-terminated with the unconditioned stimulus (US), a 5 ms periorbital air puff (2 psi) delivered to the right eye. Each daily session consisted of 100 CS–US paired trials and 10 randomly interleaved CS-only trials, with inter-trial intervals randomized between 4 and 12 seconds. Training was conducted over 8–9 days.

During trials, the right eye was recorded at 280 frames per second using a high-speed infrared camera (Mako U-029B, Allied Vision) equipped with a macro lens (1/2 inch, 4–12 mm, f/1.2, Tamron). Eyelid positions (upper and lower) were tracked using the open-source deep learning framework DeepLabCut ^102^, and the vertical distance between the eyelids was calculated to quantify eye closure. Conditioned response (CR) amplitude was defined as the average eyelid closure between 0.4 and 0.5 seconds after CS onset. All analyses were performed using custom MATLAB and Python code.

### Optokinetic Reflex (OKR) assay

OKR was performed as previously described ^71^. Before the experiment, mice were implanted with a head bracket and allowed to recover for at least five days. During testing, mice were head-fixed on a treadmill (speed, 2.5 cm/s) with the left eye positioned at the center of a custom-made apparatus. The apparatus was fully enclosed for light control. Visual stimuli were presented on a cylindrical paper screen positioned 25 cm from the left eye and generated by a laser screen beam projector (MP-CL1A, 1920 × 720; Sony, Inc.). The stimuli consisted of vertical bars (7° spacing, 3° width) that moved horizontally back and forth with a ±5° amplitude. The left eye was recorded at 200 fps (640 × 480 pixels) using a high-speed IR camera (Mako U-029B; Allied Vision, Exton, PA). Visual stimuli and camera acquisition were controlled by a custom MATLAB GUI. An infrared LED mounted above the camera generated a corneal reflection, and a second IR LED positioned below the camera illuminated the left eye. One day prior to testing, mice were habituated to the apparatus for 20 min. During experiments, pilocarpine (4% ophthalmic drops; Patterson Vet Supply, Inc.) was applied to the left eye to limit pupil dilation and thereby improve tracking accuracy in darkness.

Immediately before eye movement recording, we acquired images of each pupil while moving the camera ±10° around the vertical axis of the turntable to calculate the radius of pupil rotation (Rp), which was used to convert pixel position to eye angular position ^103^. Rp was measured at three different pupil diameters using Rp = Δ/sin(20°), where Δ is the pixel distance between pupil centers determined from their positions relative to the corneal reflection in images taken at ±10°. A regression line relating Rp to pupil diameter was then generated to estimate Rp across diameters. Angular eye movement between times t1 and t2 was computed as:

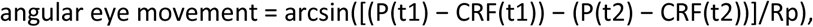

where P is the pupil center position, CRF is the corneal reflection position, and Rp is the value estimated from the regression for the corresponding pupil diameter.

For OKR measurements, the left eye was recorded while visual stimuli of different temporal frequencies (0.2, 0.5, and 1 Hz) and fixed ±5° amplitude were presented with the turntable stationary. Pupil center positions were extracted using DeepLabCut ^102^. Pixel velocities were converted to angular velocities, then low-pass filtered at twice the stimulus frequency. Data segments with eye velocities >40°/s were excluded to remove saccades. OKR gain was calculated as the ratio of the amplitude of angular eye velocity to the angular velocity of the visual stimulus. All analyses were performed using custom MATLAB and Python code.

### Open field

Mice were placed in an uncovered rectangular arena (30.3 × 45.7 cm, 30.5 cm high) and allowed to explore freely for 10 minutes while being recorded using a monochrome infrared camera (Mako U-130B, Allied Vision). Mouse position was tracked using custom MATLAB code. Total distance traveled and time spent in the center zone were calculated. The center zone was defined as the four central regions when the arena floor was evenly divided into 16 sections.

### Light-dark chamber

The light–dark apparatus (40 × 20 cm) consisted of a brightly illuminated chamber (>600 lux) and a dark chamber (<10 lux) connected by a doorway. Mice were initially placed in the dark chamber and allowed to freely explore both chambers for 10 minutes while being recorded. Position tracking was performed using custom MATLAB code, and the ratio of time spent in the light chamber was calculated.

### Elevated plus maze

Mice were placed on an elevated plus maze consisting of two open arms and two closed arms (each arm 30 x 5 cm, 50 cm high) and allowed to explore for 10 minutes. Behavior was recorded using a camera (ELP, Ailipu Technology Co.; 30 Hz, 640 × 480 pixels) with iSpy open-source software. Position tracking was performed using custom MATLAB code, and the ratio of time spent in the open arms was calculated.

### Three-chamber assay

The apparatus consisted of a clear rectangular Plexiglas box (40.5 cm wide, 60 cm long, 22 cm high) divided into three equal compartments by two transparent walls, each with a door allowing access between chambers. Mice were first habituated to the middle chamber for 5 minutes with the side doors closed. During a subsequent 10-minute baseline session, the doors were opened, allowing free exploration of all three chambers while being recorded. The mouse was then confined to the middle chamber while stimuli were introduced. For the sociability test, a wire cup (10 cm diameter) containing a social stimulus (a same-sex juvenile mouse, 30 days old) was placed in one side chamber, while an identical cup containing a novel object (mouse-sized plastic toy, Schleich GmbH, Germany) was placed in the opposite chamber. The placement of social and object stimuli was randomized. Mice were then allowed to explore all chambers for 10 minutes while being recorded. For the social novelty test, the object was replaced with a novel mouse (same sex, 30 days old, from a different cage than the first stimulus mouse), and mice were again allowed to explore for 10 minutes while being recorded. Time spent in the social, object, and novel chambers was quantified using custom MATLAB code.

## Statistical analysis

All statistical tests, significance analyses, number of individual experiments (n), and other relevant information for data comparison are specified in **Supplementary Table 1**. Statistical analysis was performed using GraphPad Prism 10. Significance levels are indicated as*p <0.05, **p <0.01, ***p < 0.001, ****p < 0.0001, and not significant (ns). Power analysis was used to predetermine sample sizes. The ROUT method (Robust Regression and Outlier Removal) was used to exclude extreme outliers. In box plots, the bottom and top of each box are the 25th and 75th percentiles of the sample, respectively. The distance between the bottom and top of each box is the interquartile range. The bottom and top error bars indicate the min and max, respectively.

## Supporting information

main supplementary information file

## Acknowledgements

We thank D. Ginty, G. Fishell, C. Chen and H. Fujita for comments on the manuscript. This work was supported by the NIH (grant nos. R35NS097284 and R35142971 to W.G.R.), the Edward R. and Anne G. Lefler Center, and Hock E. Tan and K. Lisa Yang Center for Autism Research.

