## Supplementary material for "Specialized outputs and behavioral contributions of Purkinje cell subtypes": main supplementary information file

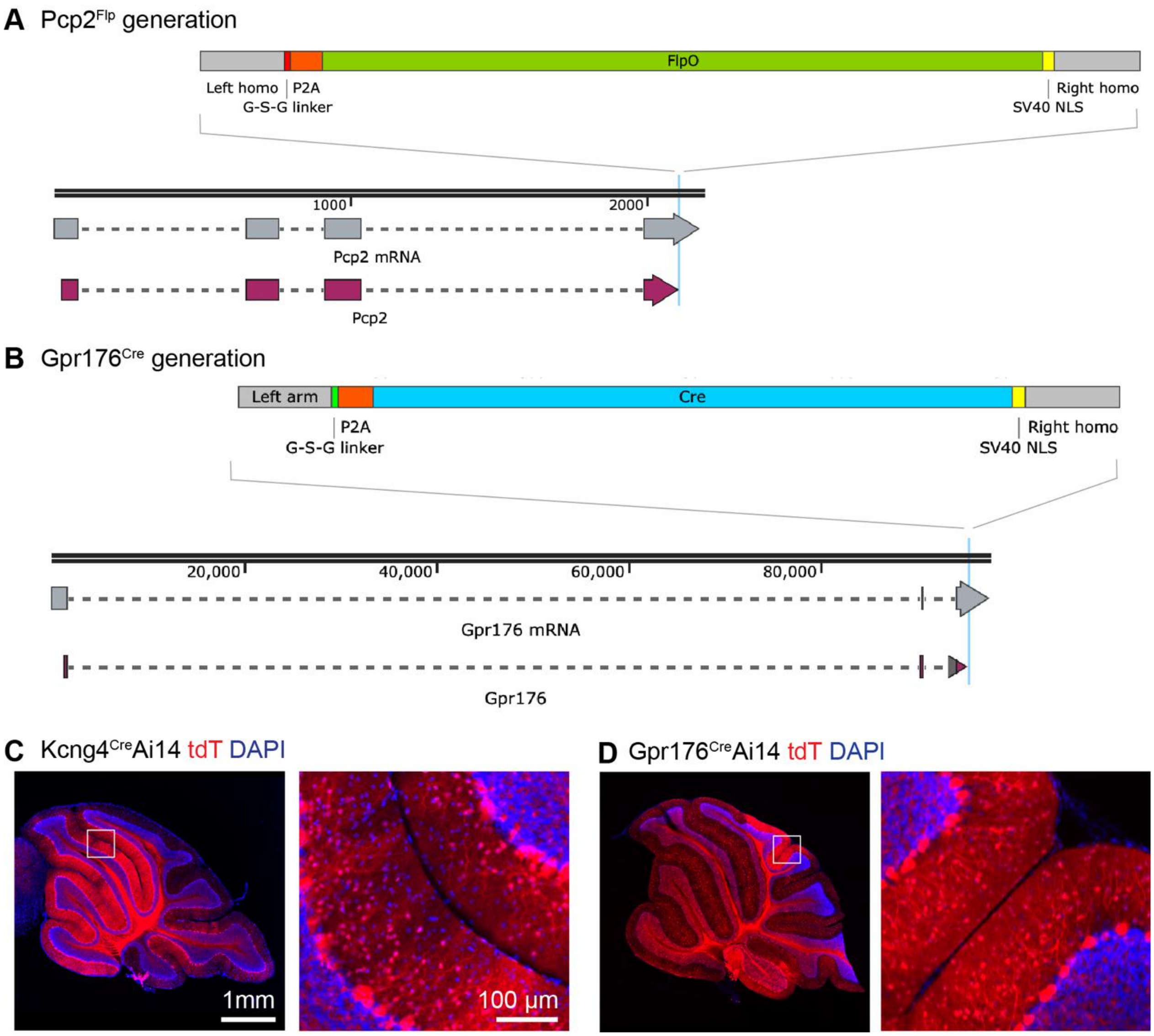


**Figure S1. Generation and validation of the mouse lines, related to Figure 1.**

**A-B.** Generation of Pcp2$^{Flp}$ and Gpr176$^{Cre}$ mice. DNA fragments containing either a Flp recombinase or a Cre recombinase were inserted before the stop codon of Pcp2 or Gpr176 without affecting the coding sequence.

**C**. A sagittal section in the Kcng4$^{Cre}$Ai14 mouse, in which Kcng4$^{Cre}$ labels both PCs and MLIs (tdT).

**D**. Same as **C** but for Gpr176$^{Cre}$Ai14 mouse, in which Gpr176$^{Cre}$ labels both PCs and MLIs (tdT).

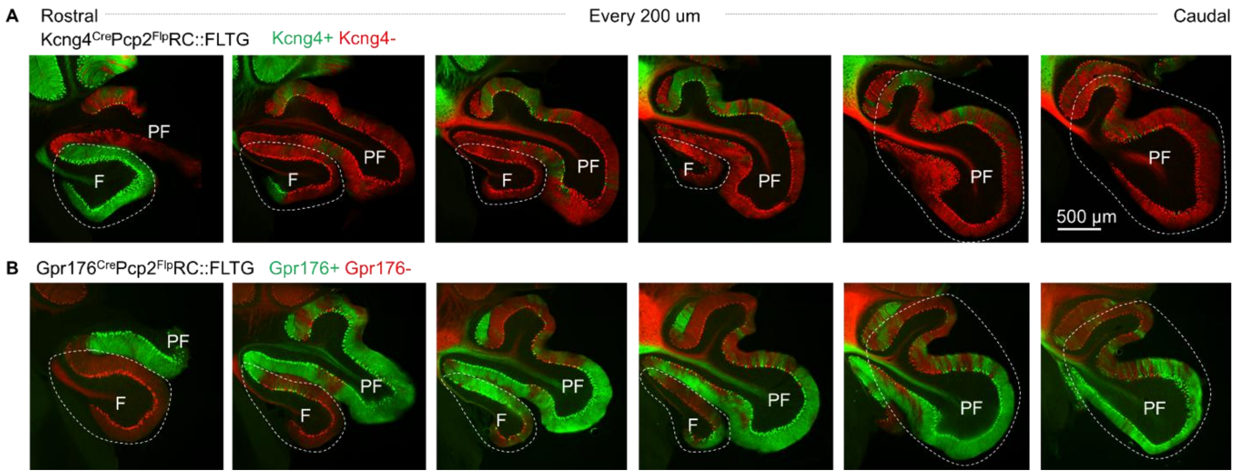


**Figure S2. Flocculus and paraflocculus signal in Kcng4$^{Cre}$Pcp2$^{Flp}$RC::FLTG and Gpr176$^{Cre}$Pcp2$^{Flp}$RC::FLTG mice, related to Figure 1.**

**A.** Serial coronal sections across the flocculus and paraflocculus in a Kcng4$^{Cre}$Pcp2$^{Flp}$RC::FLTG mouse showing the distribution of Kcng4+ (green) and Kcng4- PCs (red). Kcng4+ PCs are mainly found in rostral F.

**B.** same as **A** but for Gpr176$^{Cre}$Pcp2$^{Flp}$RC::FLTG. Gpr176+ PCs are mainly found in ventral PF.

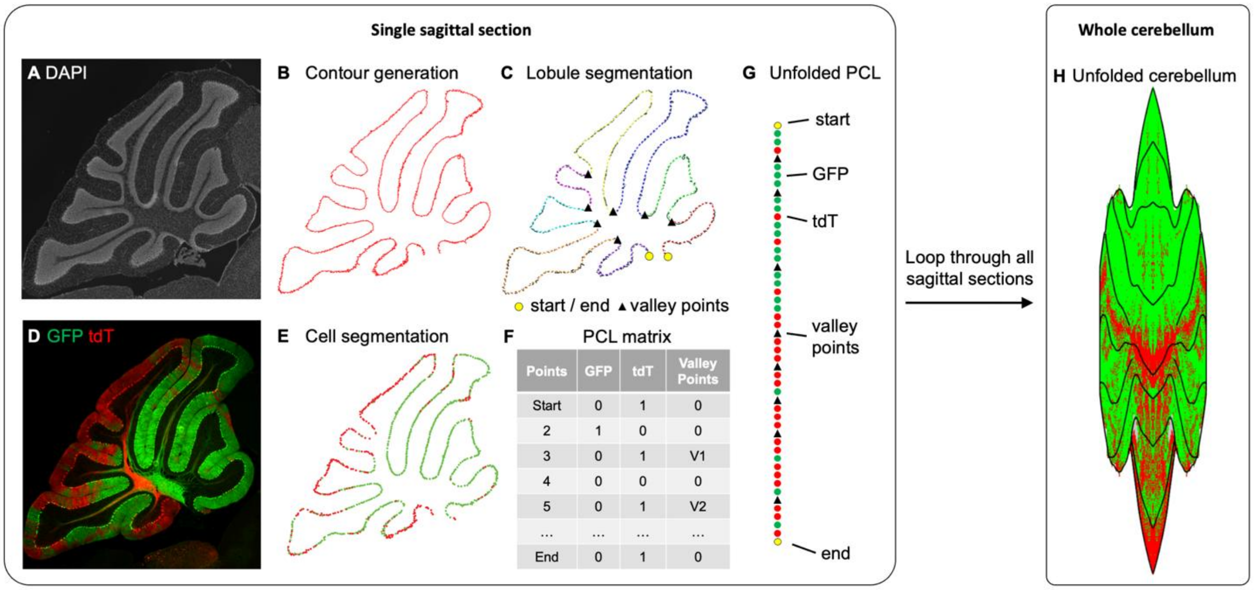


**Figure S3. Methods for unfolding the cerebellar cortex, related to Figure 2.**

**A.** DAPI signal of a sagittal section for demonstration.

**B.** Automatically generated PCL contour based on **A**.

**C.** Automatically generated smoothened contour and valley points (lobule boundary) based on **B**.

**D.** GFP and tdT signal of the same sagittal section.

**E.** Automatically segmented GFP+ and tdT+ PC cell bodies based on **D**.

**F.** Matrix representation of the PCL, with information of the length, GFP, tdT, and valley points.

**G.** Line representation of the unfolded PCL, with symbols showing start, GFP+ and tdT+ cell bodies, valley points, and end.

**H.** The same process is looped through all sagittal sections from half of the cerebellum, generating serial matrix representations, which are presented as lines and aligned to generate an unfolded map of the half cerebellar cortex. The image was mirrored to generate a full cerebellar cortex.

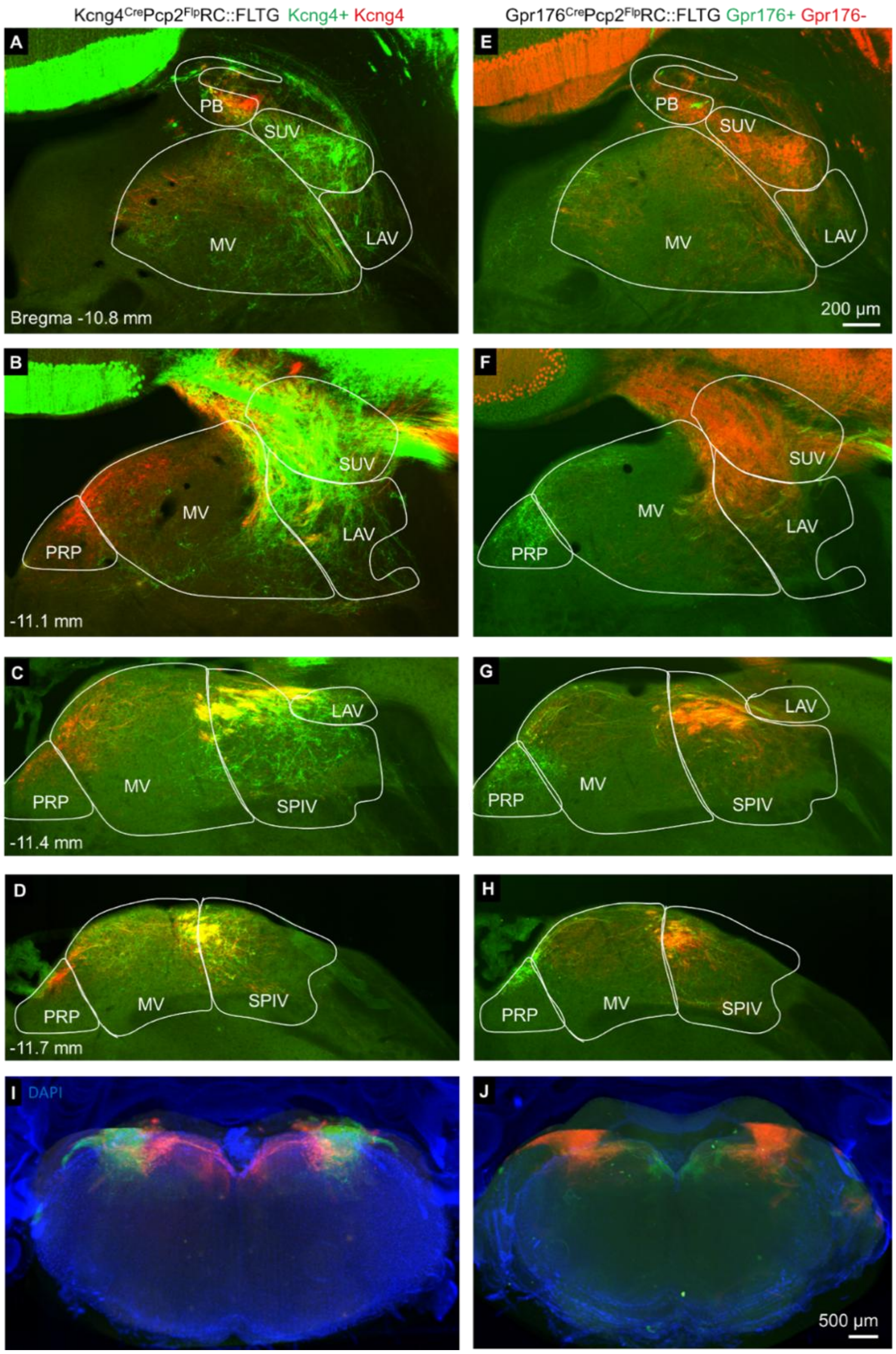


**Figure S4. Brainstem projections by PC subtypes, related to Figure 4.**

**A-D.** Serial coronal sections of the right brainstem across multiple bregma positions showing the axon projections by Kcng4+ (green) and Kcng4- (red) PCs in a Kcng4$^{Cre}$Pcp2$^{Flp}$RC::FLTG mouse. Kcng4+ PCs mainly innervate lateral brainstem, including SUV, LAV, SPIV, and ventral-lateral MV, whereas Kcng4- PCs innervate PB, PRP, and dorsal-medial MV.

**E-H**. Same as **A-D** but for a Gpr176$^{Cre}$Pcp2$^{Flp}$RC::FLTG mouse, in which Gpr176+ PC axons are labeled with GFP (green) and Gpr176- PC axons are labeled with tdT (red). Gpr176+ PCs mainly project to PRP and dorsal-medial MV, similar to Kcng4- PC but with a more restricted pattern.

**I**. Caudal view of a 3D reconstruction of the brain showing the axon projections to the brainstem by Kcng4+ (green) and Kcng4- (red) PCs in a Kcng4$^{Cre}$Pcp2$^{Flp}$RC::FLTG mouse. Signal outside of brainstem were masked out.

**J**. Same as **I** but for a Gpr176$^{Cre}$Pcp2$^{Flp}$RC::FLTG mouse.

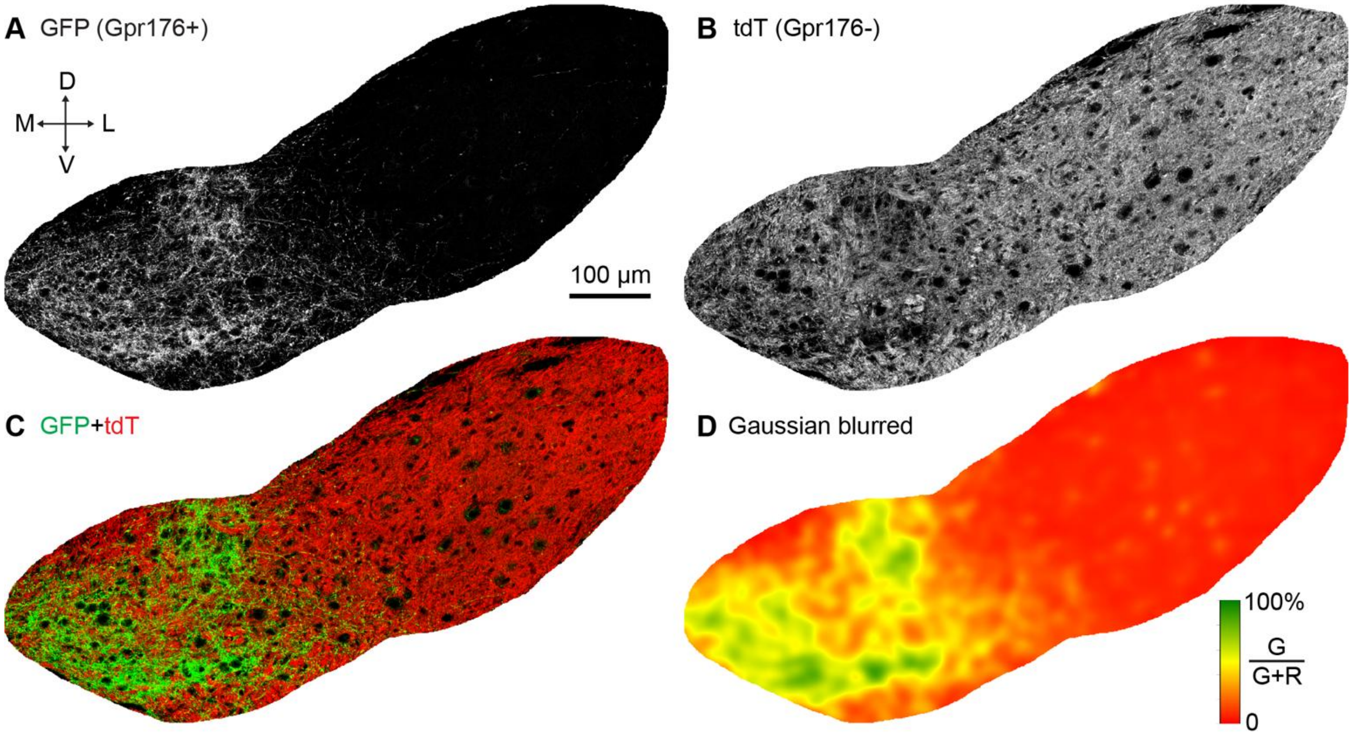


**Figure S5. Confocal and Gaussian-blurred images for a FN section of a Gpr176$^{Cre}$Pcp2$^{Flp}$RC::FLTG mice, related to Figure 5.**

**A.** Greyscale confocal image of GFP labelling of Gpr176+ PCs axons and boutons in the FN (single Z plane).

**B.** As in **A** but for tdT labelling of Gpr176- PCs.

**C.** Overlay of GFP (*green*, Gpr176+ PCs) and tdT labeling (*red*, Gpr176- PCs).

**D.** Gaussian blur with the ratio of GFP to tdT labeling shown (based on a confocal stack).

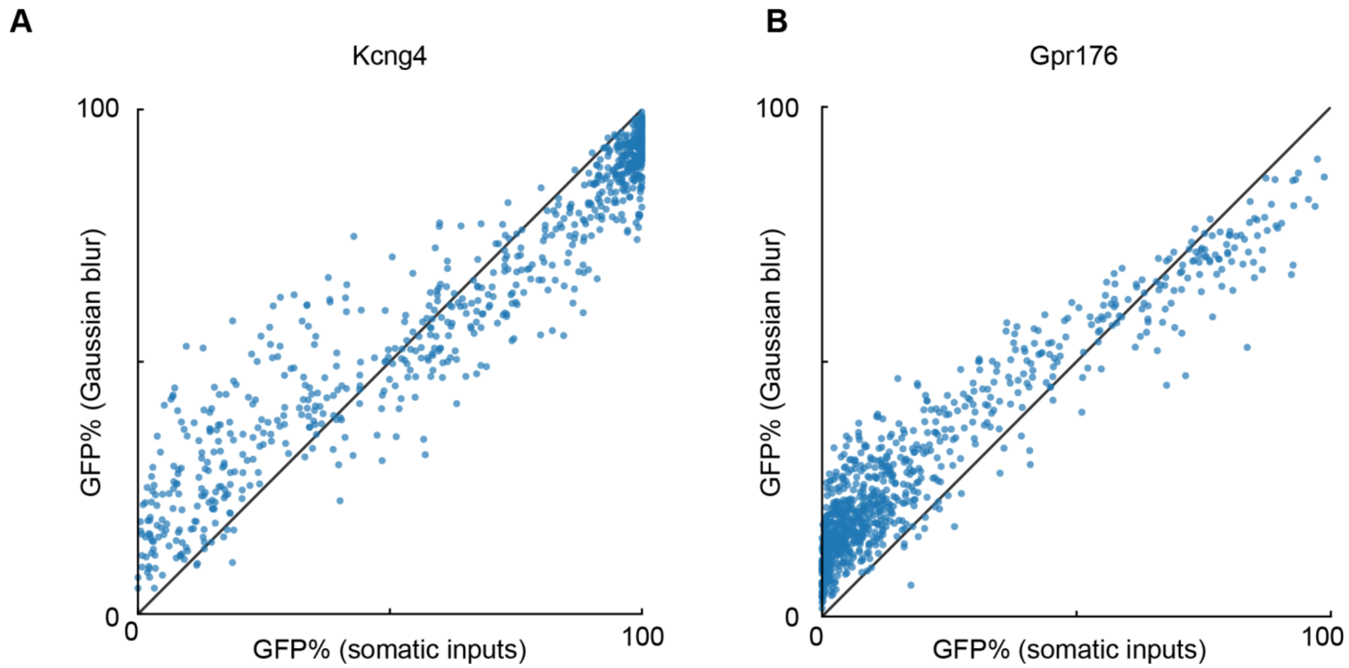


**Figure S6. Comparison of G/(G+R) ratio in Gaussian-blurred images and soma input quantification, related to Figure 5.**

**A.** G/(G+R) ratio obtained at the centroid of each FN neuron in Gaussian-blurred image vs. the ratio quantified based on inputs to the soma of each FN neuron in Kcng4$^{Cre}$Pcp2$^{Flp}$RC::FLTG mice.

**B.** Same comparison but for Gpr176$^{Cre}$Pcp2$^{Flp}$RC::FLTG mice.

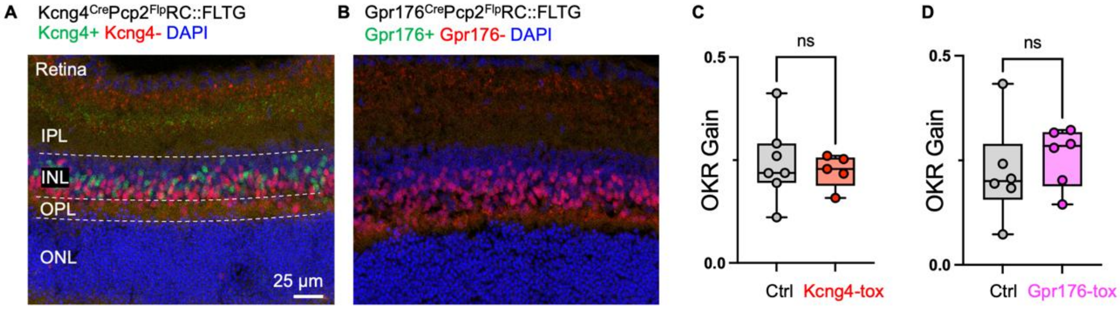


**Figure S7. Retina labeling by Kcng4$^{Cre}$ and Gpr176$^{Cre}$, related to Figure 7.**

**A.** A retina section showing subpopulations of bipolar neurons labeled as green (Kcng4+) or red (Kcng4-) in a Kcng4$^{Cre}$Pcp2$^{Flp}$RC::FLTG mouse. Note that these Kcng4+ bipolar neurons will be targeted in the Kcng4-tox mouse. IPL, inner plexiform layer; INL, inner nuclear layer; OPL, outer plexiform layer; ONL, outer nuclear layer.

**B.** Same as **A** but for a Gpr176$^{Cre}$Pcp2$^{Flp}$RC::FLTG mouse showing subpopulations of bipolar neurons labeled as red (Gpr176-Pcp2+). No Gpr176+Pcp2+ cells (green) are detected in the retina, therefore bipolar neurons are not affected in the Gpr176-tox mice.

**C.** Optokinetic reflex assay (OKR) was performed in ctrl and Kcng4-tox mice to examine the visual functions, in which visual stimulation (vertical bars moving left and right) is presented at 0.5 Hz to a head fixed mouse. OKR gain is defined by (pupil velocity)/(peak velocity of visual stimulation). There is no difference in the average OKR gain between ctrl (n=7) and Kcng4-tox (n=5) mice.

**D.** Same as C but for ctrl and Gpr176-tox mice. There is no difference in the average OKR gain between ctrl (n=6) and Gpr176-tox (n=6) mice.

**Supplemental video 1**. A 3D reconstruction of the cerebellum in a $Kcng4^{Cre}Pcp2^{Flp}$RC::FLTG mouse showing the distribution of Kcng4+ (green) and Kcng4- (red) PCs in the cerebellar cortex, related to Figure 1.

**Supplemental video 2**. A 3D reconstruction of the cerebellum in a $Gpr176^{Cre}Pcp2^{Flp}$RC::FLTG mouse showing the distribution of Gpr176+ (green) and Gpr176- (red) PCs in the cerebellar cortex, related to Figure 1.

**Supplemental video 3**. A 3D reconstruction of the cerebellum in a $Kcng4^{Cre}Pcp2^{Flp}$RC::FLTG mouse showing the axon projections by Kcng4+ (green) and Kcng4- (red) PCs in the CbN. Signals outside of CbN were masked out, related to Figure 4.

**Supplemental video 4**. A 3D reconstruction of the cerebellum in a $Gpr176^{Cre}Pcp2^{Flp}$RC::FLTG mouse showing the axon projections by Gpr176+ (green) and Gpr176- (red) PCs in the CbN. Signals outside of CbN were masked out, related to Figure 4.

**Supplemental video 5**. A 3D reconstruction of the brain in a $Kcng4^{Cre}Pcp2^{Flp}$RC::FLTG mouse showing the axon projections by Kcng4+ (green) and Kcng4- (red) PCs in the brainstem. Signals outside of brainstem were masked out, related to Figure 4.

**Supplemental video 6**. A 3D reconstruction of the cerebellum in a $Gpr176^{Cre}Pcp2^{Flp}$RC::FLTG mouse showing the axon projections by Gpr176+ (green) and Gpr176- (red) PCs in the brainstem. Signals outside of brainstem were masked out, related to Figure 4.

**Supplemental video 7.** A representative gait video in ctrl (*top*) and Kcng4-tox (*bottom*) mice showing different body parts extracted from bottom view videos and side view videos, related to Figure 7.

**Supplemental video 8.** A representative gait video in ctrl (*top*) and Gpr176-tox (*bottom*) mice showing different body parts extracted from bottom view videos and side view videos, related to Figure 8.
